# Inferring virtual cell environments using multi-agent reinforcement learning

**DOI:** 10.1101/2025.11.21.689815

**Authors:** Noah Cohen Kalafut, Chenfeng He, Jie Sheng, Pramod Bharadwaj Chandrashekar, Jerome Choi, Daifeng Wang

## Abstract

Single cells interact continuously to form a cell environment that drives key biological processes. Cells and cell environments are highly dynamic across time and space, fundamentally governed by molecular mechanisms, such as gene expression. Recent sequencing techniques measure single-cell-level gene expression under specific conditions, either temporally or spatially. Using these datasets, emerging works, such as virtual cells, can learn biologically useful representations of individual cells. However, these representations are typically static and overlook the underlying cell environment and its dynamics. To address this, we developed CellTRIP, a multi-agent reinforcement learning method that infers a virtual cell environment to simulate the cell dynamics and interactions underlying given single-cell data. Specifically, cells are modeled as individual agents with dynamic interactions, which can be learned through self-attention mechanisms via reinforcement learning. CellTRIP also applies novel truncated reward bootstrapping and adaptive input rescaling to stabilize training. We can in-silico manipulate any combination of cells and genes in our learned virtual cell environment, predict spatial and/or temporal cell changes, and prioritize corresponding genes at the single-cell level. We applied and benchmarked CellTRIP on various simulated and real gene expression datasets, including recapitulating cellular dynamic processes simulated by gene regulatory networks and stochastic models, imputing spatial organization of mouse cortical cells, predicting developmental gene expression changes after drug treatment in cancer cells, and spatiotemporal reconstruction of Drosophila embryonic development, demonstrating its outperformance and broad applicability. Interactive manipulation of those virtual cell environments, including in-silico perturbation, can prioritize spatial and developmental genes for single-cell-level changes, enabling the generation of new insights into cell dynamics over time and space. CellTRIP is open source as a general tool and available at github.com/daifengwanglab/CellTRIP.

## 2 Introduction

Single cells continually interact and establish a cell environment to coordinate many key biological processes, such as development and disease progression. For example, within tissue microenvironments, cells develop along different lineages and migrate to specific regions during maturation of cell types, such as T cell lineages^1^. Furthermore, gene expression, a key single-cell-level mechanism to determine cellular function, can be controlled by intracellular gene regulation and through intercellular molecular interactions with neighboring cells^2,3^. These interactions are dynamic, developing the cell environment to drive specific biological processes. These environments are sensitive to small perturbations. Varying conditions can lead to heterogeneous functional outcomes. For example, inter-individual variance in tumor immune microenvironments can dictate responsiveness to chemo-immunotherapy^4^. Characterizing these environments is critical to understanding a wide range of developmental, pathological, and embryonic processes, among others.

Next-generation sequencing techniques enable measurement of genome-wide molecular activities, including gene expression, especially at the single-cell level^5^. Many of those datasets can also provide dynamic information, including spatial positioning^6^ and temporal annotation ^7,8^. Typically, these sequencing-based measurements are limited to one or few time points over the course of a particular biological process. This temporal scarcity challenges a deeper understanding of dynamic interactions within the cell environment governing many biological processes, including developmental trajectories and perturbation outcomes. Some recent single-cell datasets can contain time-dependent information, such as for cell maturity and disease progression. Thus, many trajectory inference methods have been developed to computationally learn dynamic patterns of single cells using these datasets. For example, MOSCOT^9^ uses optimal transport across multimodalities to recover cell developmental trajectories from time-series and spatial data. Other methods like URD^10^ and STREAM^11^ recover gene expression changes along cell developmental timepoints. None of these methods, however, focus on modeling underlying dynamic cell-cell interactions and cellular behaviors, and are therefore unable to characterize the cell environment.

Reinforcement learning (RL), a machine learning approach for modeling agent-environment interactions and discovering optimal strategies through interaction, has been widely applied across many fields, including robotics^12,13^, DNA and protein sequence design^14^, and manufacturing^15^. Broadly speaking, RL seeks to train decision-making agents to optimize a reward function through interaction with a given environment. This optimization is performed through repeated trial-and-error, reinforcing behaviors that lead to positive outcomes. One of the most prevalent and flexible RL approaches is Proximal Policy Optimization (PPO). PPO was developed as an efficient objective function and training procedure that uses several user-adjustable parameters to efficiently reinforce model behaviors that increase a reward value^16^. RL can also model a typical cellular process. In particular, cells can be represented as agents that interact in an unseen cell environment. For example, reinforcement learning has been applied to modeling cell movement and division and validated related pathways during embryonic development of *C elegans* ^17^. Some RL-based methods have started to analyze single-cell data, including scRL^18^, which uses an actor-critic architecture to predict cell fate under disease, development, and gene-knockdown conditions. CNRein^19^ also uses single-cell data to infer realistic copy number aberrations by inferring a cancer evolutionary model for individual patients. However, neither method models the underlying cell environment or incorporates dynamic information. Despite its potential, RL has not been widely applied to general analysis of emerging large-scale single-cell data, which is readily available and can provide useful dynamic information from the cell environment under many temporal and spatial conditions.

To address this, we developed CellTRIP, a multi-agent reinforcement learning method to infer a scalable and interactive virtual cell environment that can be manipulated at the gene or cell level. CellTRIP is able to perform spatial or temporal imputation, predict in-silico perturbation using single-cell data in both time and space domains, and recover missing cell developmental stages. CellTRIP utilizes PPO combined with generalized advantage estimation^20^, PopArt^21^, and well-known optimizations^22,23^. We show that CellTRIP (1) outperforms existing methods for recovering developmental trajectories on simulation data^24^, (2) imputes spatial transcriptomics data from the mouse cortex^25^, (3) predicts the effects of gene knockdown on drug perturbation datasets from cancer cell lines^7^, and (4) recovers spatiotemporal cell organization of developmental stages and infers development and tissue-specific genes during *Drosophila* embryonic and larval development^8^.

## 3 Methods

CellTRIP infers a *virtual cell environment* from the single-cell data (e.g., gene expression) for spatiotemporal imputation and perturbation prediction. Given a single-cell dataset, each Cell *i* has *K* types of measurements, such as gene expression and spatial location, where |*K*| is a set of indices representing each type of measurement. Let 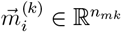 represent measured data for Cell *i* in modality *k* ∈ *K* with *n*_*mk*_ features. We also define the matrix 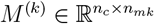, where each row is 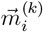 over all *n*_*c*_ cells in the dataset.

CellTRIP consists of three interconnected components that are iteratively updated during learning and inference: The environment space, environmental reward, and residual self-attention policy. In CellTRIP (Figure 1), single cells are represented as independent agents within an *environment space*. The environment space is a coordinate space of reduced dimensionality, defining cell positions and velocities. Cell positions, velocities, and single-cell data are supplied to the *residual self-attention policy*, which computes cell actions (i.e., changes in velocity) for all cells simultaneously. Cells use such actions to interact with and navigate through the environment space. CellTRIP agents try to maximize the *environmental reward*, which increases as the imputation loss inferred from the environment space (cell positions and velocities) decreases. An additional pinning module is utilized to impute or align single-cell modalities, including spatial coordinates. Based on the optimized virtual cell environment, we can perform spatiotemporal imputation and predict perturbation outcomes.

**Figure 1.**
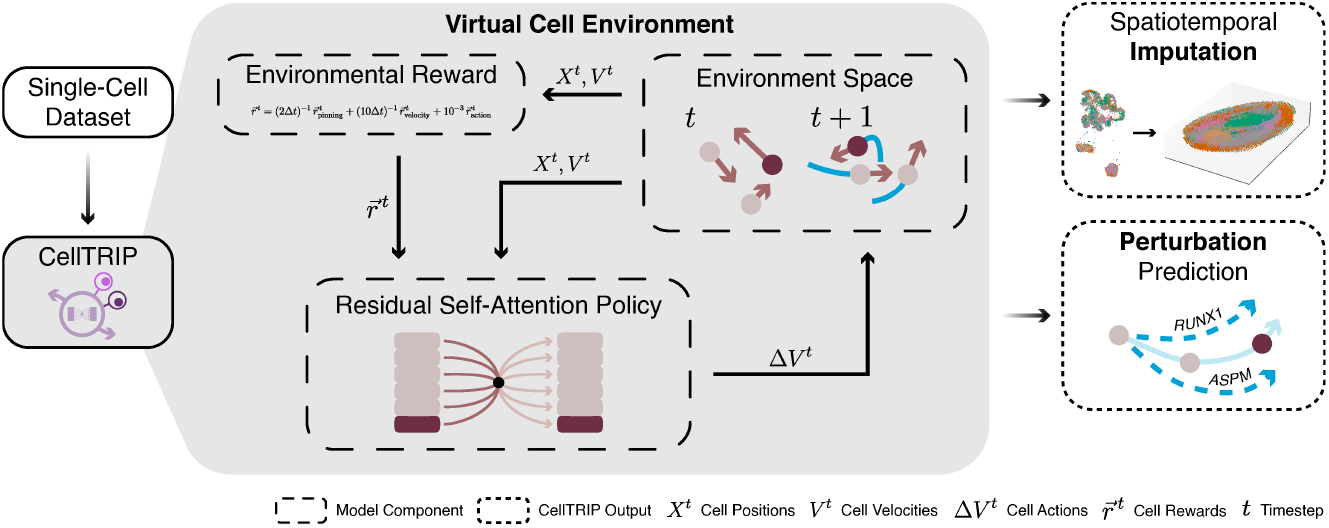
CellTRIP uses multi-agent reinforcement learning techniques to infer virtual cell environments from static biological data for spatiotemporal imputation and perturbation prediction. The model learns to optimally construct environment spaces to maximize data retention from source modalities. Retention quality and reliability are communicated to the model through environmental rewards, while the model can interact with the environment by imparting forces on each cell, independently. A residual selfattention module is utilized cell and neighbor embeddings to determine the velocity changes of each cell from one timestep to another. After training, the model can be used in various environments for spatiotemporal imputation, to simulate cell development, or for perturbation prediction.

### 3.1 Environment Space

The environment space calculates cell positions and velocities, simulating cell state changes over time. Each simulation runs for a user-specified amount of time, 128 seconds by default, with each timestep t representing Δ*t* seconds of simulation time, defaulting to 0.1s (totaling 1,280 timesteps). During training, the total simulation time is uniformly sampled between 64 and 128 seconds (See Supplementary Section S6). At a particular timestep t, each Cell i has a cell position 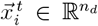 and cell velocity 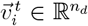, where n_*d*_ is the user-specified dimensionality of the environment space (32 by default). Cell positions and velocities of all given cells may also be represented as matrices 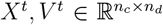, where row i corresponds to 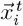 or 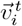, respectively. Aggregating cell positions, velocities, and single-cell data, we get the cell states 𝒮^*t*^ = {*X*^*t*^, *V*^*t*^} + {*M*^(*k*)^ for *k* ∈ *K*_sources_} . The environment also stores a list of source and target modalities, *K*_sources_, *K*_targets_ ⊆ *K*, which determine the policy input and predicted modalities, respectively (Supplementary Section S5.1). For instance, we can use single-cell gene expression data (source modality) to predict spatial locations (target modality).

At each timestep t, the environment space gives a state matrix 𝒮^*t*^ to the policy (Section 3.3), which returns a cell action matrix Δ*V*^*t*^. The cell velocities and positions are then updated according to the following rules:

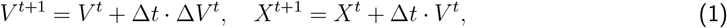

If trained correctly, the cell positions typically converge after a certain number of timesteps, i.e., 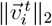 is small. We refer to the converged cell positions as the *steady state*. For a visual representation of this process, see Supplementary Figure S2.

### 3.2 Environmental Reward

The goal of the environmental reward is to maximize the ability of the policy to impute target modalities from the environment space (Supplementary Figure S3). This is accomplished through the combination of three rewards: the pinning reward, velocity penalty, and action penalty. We provide minimal guidance during training to allow the model to infer the inter-cell relationships of the input data, rather than relying on modality-specific prior knowledge.

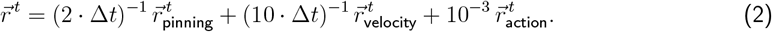

The default coefficients for each reward are shown in Equation 2, but can be tuned by users. Note that the scales of 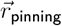 and 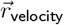 are dependent on Δ*t*. So, their coefficients are divided by Δt to remove this dependence.

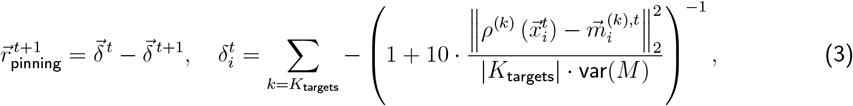

The central objective is the pinning reward, which increases as the policy-imputed modalities approach the target modalities. where *ρ*^(*k*)^ is the pinning module (Section 3.4.1) for modality k, var(·) is the variance across all matrix values, and 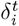 is the element of 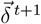 corresponding to Cell i. The result is then a weighted sum of the inverted mean squared error (MSE) between the imputed and target modalities. Observe that, as the MSE rises or drops, 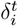 approaches 0 or −1, respectively. Also note that the result is invariant to the number of features, ensuring equitable weighting of multiple modalities.

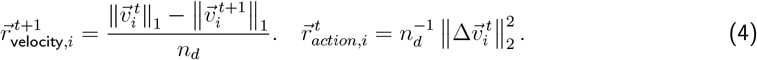

The velocity penalty is added for stability. This penalty increases with velocity in the environment space, incentivizing slower movement and disincentivizing unbounded exploration.

The pinning reward and velocity penalties reflect changes in the pinning MSE and velocity, respectively. This leads to the properties 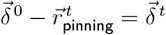 and 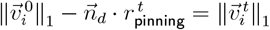. The cumulative reward across a whole simulation then reflects the corresponding cumulative change in pinning accuracy and velocity and, crucially, is not dependent on the number of timesteps.

Finally, the action penalty is used to incentivize smooth changes in velocity. Erratic movements are more difficult to predict and coordinate between agents.

### 3.3 Residual Self-Attention Policy

The residual self-attention policy is the decision-making component of CellTRIP, responsible for evaluating cell states from the environment space and predicting optimal cell actions to iteratively maximize the resultant environmental reward. Policy inference consists of several steps, including cell and neighbor embedding, cell summary embedding, and action sampling. The cell and neighbor embeddings encode cell positions, velocities, and single-cell modalities in one unified representation. The cell summary embeddings add further environmental context to each cell embedding by computing residual self-attentions between the cell and neighbor embeddings. Finally, cell actions are sampled from the cell summary embeddings to be given to the environment space.

During forward computation, for each Cell *i*, cell positions and velocities (i.e., 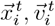 totaling 2n_*d*_ features) are concatenated with the corresponding cell information for modalities k K, resulting in a vector of size 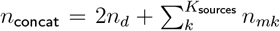. Cell and neighbor embeddings 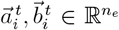 are then computed using multilayer perceptrons (MLPs) of dimension *n*_concat_ × *n*_hidden_ ×*n*_hidden_ with a Parametric ReLU (PReLU) activation function between each layer, where n_hidden_ is a user-defined parameter. We call these MLPs *E*_a_ and *E*_b_ for cell and neighbor encoders, respectively

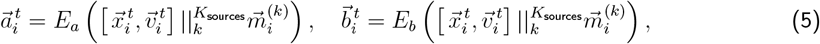

where || denotes vector concatenation. CellTRIP contains an additional parameter to generate neighbor embeddings relative to each cell by concatenating 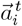 to each neighbor input vector, as in Baker et al. ^26^ . However, this increases the complexity of the forward computation by a factor of *n*_*c*_.

Sets of cell and neighbor embeddings are then run through a residual self-attention model individually. In particular, the cell embeddings (queries) and neighbor embeddings (key and value) are layer-normalized and fed through a multiheaded attention layer. The number of neighbors per cell is user-adjustable and defaults to 1k. The unnormalized cell embeddings are added to the attention output, implementing the residual component. The layer-normalized intermediate output is then fed through an additional residual two-layer MLP of hidden layer sizes 4*n*_hidden_ × *n*_hidden_ × 4*n*_hidden_ with PReLU activation and added to the unnormalized input to the MLP, generating the cell summary embeddings, 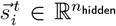. The cell summary embeddings may also be represented as a matrix 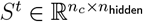 . CellTRIP contains an option to stack multiple of these blocks in sequence, but the default model only uses one.

The cell summary embeddings are fed through PReLU activation and the decider module, an MLP *E*_*s*_, with layer sizes *n*_*d*_ × 2*n*_*d*_ × *n*_*d*_ with intermediate PReLU activations followed by Tanh. The output, *E*_*s*_(*S*^*t*^), determines the final cell actions. The outputs are used as means in normal distributions with trainable variance σ for each dimension in the environment space. This is equivalent to the multivariate normal distribution, 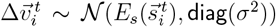, where 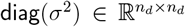 is a diagonal covariance matrix with all nonzero entries σ. Note that, internally, σ is stored in log-form to prevent negative values for σ from backpropagation. The computed cell actions are then applied to the environment space according to Equation 1 to iterate the cell positions and velocities. The model is trained using proximal policy optimization (PPO), simulating multiple environments in parallel and iterating at fixed step intervals until convergence. We additionally utilize novel input scaling techniques derived from PopArt normalization^21^. For more details concerning model training or the distributed environment, see Supplementary Section S1 and Supplementary Section S6.

### 3.4 Cell Spatiotemporal Imputation and Perturbation Prediction

With the trained CellTRIP model, specifically 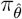 and *ρ*^(*k*)^, we can perform spatiotemporal imputation and perturbation prediction. Spatiotemporal imputation can be viewed as two major components, those being spatial and temporal imputation. CellTRIP uses pinning modules, *ρ*^(*k*)^, to estimate target modalities from the environment space, *X*^*t*^ (Section 3.4.1). These pinning modules also enable imputation of single-cell spatial coordinates from gene expression data. Temporal imputation can recover intermediate timepoints or stages from developmental data through interpolation in the environment space using the CellTRIP model, with optimal generation of pseudocells. This enables imputation of intermediate developmental stages from embryonic development time-series data. Perturbation prediction can also use the predictive capabilities of CellTRIP to predict perturbation outcomes, including gene expression, and to prioritize genes across dataset conditions. For example, if we define a perturbation function *ψ* to replace the expression of gene *g* with 0, we can prioritize larval developmental genes by anatomical region using time-series *Drosophila* data.

#### 3.4.1 Spatial Imputation

CellTRIP relies on pinning modules, *ρ*^(*k*)^ to impute from the environment space to M^(*k*)^ space, for *k* ∈ *K*_targets_. Each pinning module is an MLP with dimension *n*_*d*_ × 2*n*_*d*_ × *n*_*mk*_ with PReLU activations between layers, trained concurrently with the policy. Each pinning module uses MSE between the predicted and imputed environment spaces as its loss function.

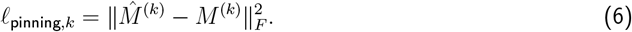

In non-spatial modalities, 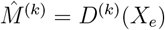, where *X*_*e*_ is the matrix of environment cell positions at a user-defined timestep (typically 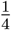 through training) in each simulation. When working with spatial data, or any data whose primary significance lies in inter-cell distances, we instead solve for a rotation matrix *R* and translation vector 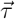 minimizing the MSE between *ρ*^(*k*)^(*X*_*e*_) and 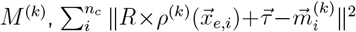 and set 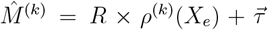. For additional details on solving for *R* and 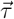 using SVD, see Supplementary Section S3.

#### 3.4.2 Temporal Imputation

We can also use CellTRIP to impute intermediate time points and cell types, even with unmatched cells. If dataset timepoints do not contain matched cells, we can use optimal transport^27,28^to estimate correspondence between the observed initial and terminal stages in the environment or imputed space. Specifically, we solve

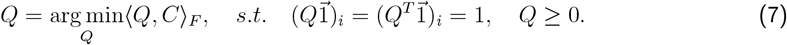

where *n*_*c*1_ and *n*_*c*2_ are the numbers of cells in the initial and terminal stages, respectively, and 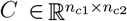 is the euclidean distance matrix between the (properly rotated) cells in the two stages. Once we have the transition matrix *Q*, we compute pseudocells by taking the mean of cells with nonzero entries in each row of *Q*. Then, we have a 1 : 1 mapping of cells in the initial and terminal stages. In practice, we preempted this process by generating pseudocells using K-means in each stage.

We then begin simulation of the initial stage pseudocells. Once the steady state is reached, we replace the input modalities *M*^(*k*)^ for *k ∈ K*_source_ with those of the terminal stage pseudocells. Simulating to steady state provides a transition tensor of dimension [timesteps × pseudocells × positions]. We can then manually choose a timestep as an intermediate timepoint. This process is outlined in Figure S1.

#### 3.4.3 Perturbation Prediction

In general, CellTRIP feature perturbation revolves around modification of *M*^(*k*)^, *k* ∈ *K*_sources_. We notate this modification through a perturbation function *ϕ*( {*M*^(*k*)^ for *k ∈ K*_sources_} ). Any number of feature perturbation techniques can be applied to CellTRIP, but we primarily focus on feature knockdown. In particular, we define 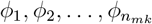, each replacing the corresponding modal feature in a chosen modality k with 0. In the case of gene expression features, starting from our steady state position and velocity, 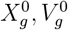, calculated without perturbation, we can simulate each perturbation function *ϕ*_*g*_,

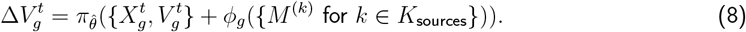

The ending positions at time T are denoted 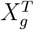. CellTRIP also contains additional knockdown strategies, which can be found in Supplementary Section S7.1. The transition states between 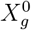 and 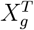 are then computed to quantify the *gene effect size* and *gene trajectory length* for each Cell *i*,

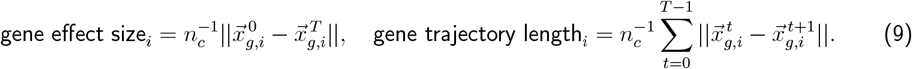

We may also compute the gene effect size and gene trajectory length after imputation using the pinning module, if desired. For example, we can calculate gene effect size in environment or gene expression space.

## 4 Results

### 4.1 Recapitulating Cellular Dynamic Processes Simulated by Gene Regulatory Networks and Stochastic Models

To showcase the ability of CellTRIP to recover cell state developmental genes and trajectories, we generated 1,500 simulation cells with 2,400 gene and protein expression features each using Dyngen^24^. The data consists of cells at seven distinct stages in development (Figure 2a). Possible developmental paths bifurcate twice, ending at one of three terminal cell states.

**Figure 2.**
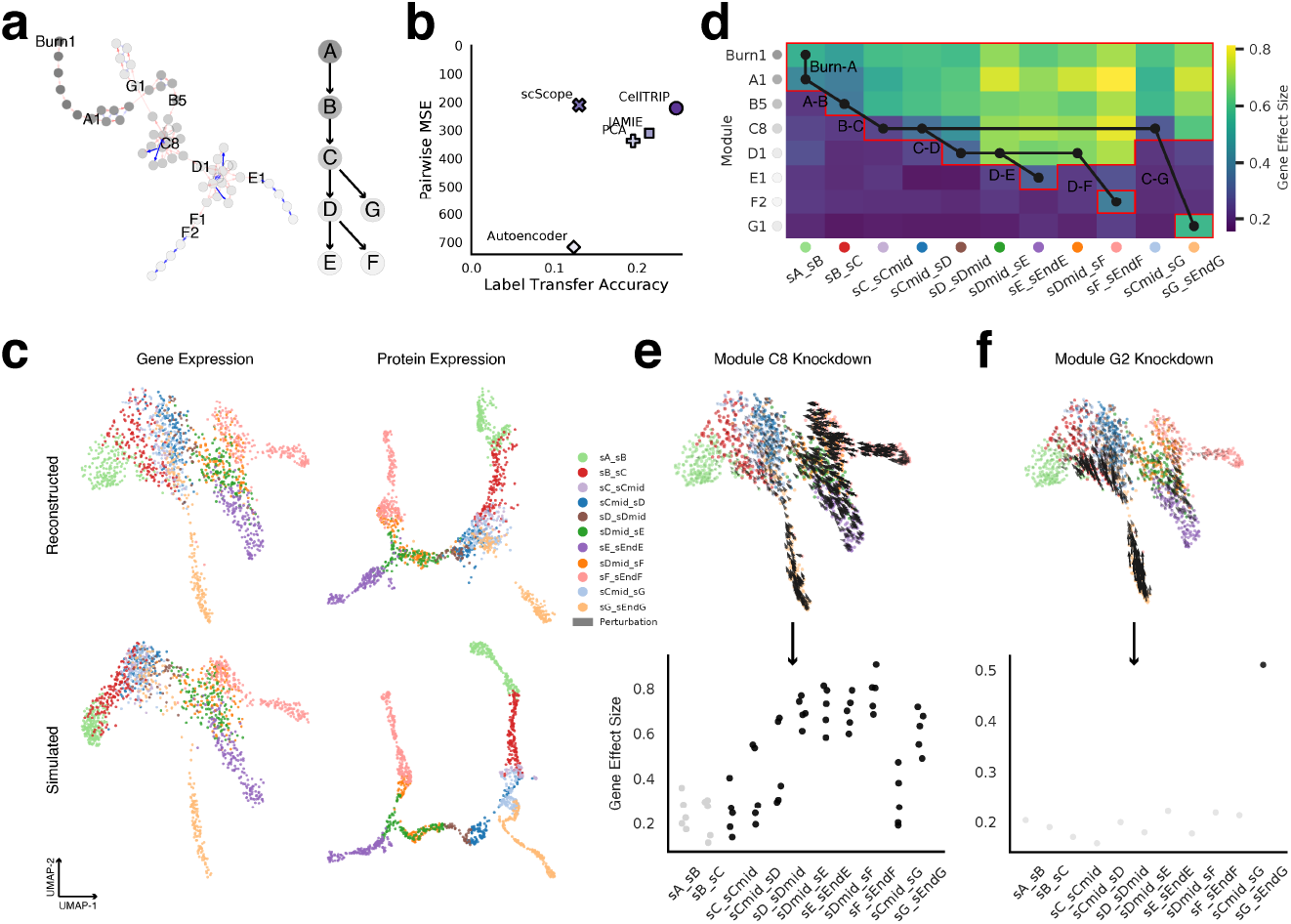
Gene prioritization and knockdown on simulation data generated from gene regulatory networks and stochastic models^24^. **a.** Ground-truth developmental module progression (Left) and cell state developmental tree (Right), shaded by cell state. Red and blue arrows indicate positive and negative regulatory effects, respectively. **b**. Label transfer accuracy (x-axis) and pairwise MSE (y-axis) for CellTRIP and comparable method reconstructions of gene expression, limited to 32 dimensions (See Section S2.1). **c**. UMAP of CellTRIP-reconstructed (Top) and ground-truth simulated (bottom) gene expression (Left) and protein expression (Right), colored by cell trajectory. **d**. Heatmap of mean CellTRIP-predicted gene effect sizes in environment space by cell trajectory (x-axis) and module (y-axis). Dots and lines indicate ground-truth developmental paths from (a), while red outlines indicate expected regions of effect for module knockdown. **e**. CellTRIP-predicted gene expression trajectory (Left) and gene effect size (Bottom) under module C8 knockdown and knockdown of constituent transcription factors, respectively (See Section 3.4.3). **f**. Panel (e), repeated for module `G2`.

We began by examining the per-modality reconstructions from CellTRIP, which were predicted from a common environment space after it reached steady state. Comparing CellTRIP reconstructions to similar methods with the same dimensionality, we observed that CellTRIP has the second lowest MSE of ∼221, closely behind scScope at ∼211, and the highest label-transfer accuracy (0.248, Figure 2b). We see that cell trajectories are distinct in the CellTRIP-reconstructed UMAPs and match the ground-truth simulation data closely (Figure 2c). In particular, cell state lineages match the ground-truth developmental tree.

After accurately reconstructing gene and protein expression modalities, we performed in-silico knockout of individual TFs per module, finding that we were able to accurately match expected gene effect size trends from the ground truth data (Figure 2d). Specifically, we chose one module from each cell state and used the mean environment space gene effect sizes from the TF knockdowns to generate a heatmap of module downstream effects across trajectories and cell states. We observed greater gene effect sizes in cells at later stages of development. Moreover, the magnitude of the gene effect sizes tended to increase with the lineage distance between cell states. The resultant gene effect size patterns also matched the expected trends from the ground-truth data, i.e., that child cell states were the most affected. Similar patterns are also seen across all modules (Supplementary Figure S5).

Looking into module C8 in particular, performing knockdown of all TFs simultaneously caused large expression changes in differentiated cell states (Figure 2e). Interestingly, some terminal cell states were less affected towards the end of their development. Namely, terminal cell states originating from cell state D (i.e., Cell states E and F), itself a child of cell state C, seemingly exhibit gene expression changes of lower magnitude than those with closer lineage to cell state C (i.e., cell state G). Looking at individual TF knockdowns for module C8, we observe the gene effect size increasing as the cells developed further from cell state C, indicating a deviating developmental trajectory caused by TF knockdown. Knockdown of modules corresponding to a terminal state results in the expected isolated effect, as can be seen from the knockdown of modules G2 or F1, consisting of only one TF each (Figure 2f, Supplementary Figure S4).

### 4.2 Imputing Spatial Organization of Mouse Cortical Cells

To demonstrate the ability of CellTRIP to impute single-cell spatial coordinates from gene expression, we trained a CellTRIP model on Visium gene expression data from 1,075 spatially-resolved cells from the adult mouse frontal cortex across six cortical layers ^29,30^. As independent validation, we imputed the spatial coordinates of 4,785 cells from an independent scRNA-seq dataset in the adult mouse frontal cortex^25^. We also used smFISH data, consisting of 2,360 cells, from the primary visual cortex^33,34^ to provide additional per-layer reference cell type proportions.

We compared several mapping-based spot assignment methods with CellTRIP. We note that CellTRIP has several key distinctions from these methods, including being a regression model which computes spatial coordinates directly from gene expression, rather than choosing from a list of known spot positions. CellTREK^30^ and Cytospace^32^ also have additional cell filtering methodologies, with CellTREK predicting 3,899 possible coordinates for 2,347 validation cells and Cytospace predicting assigning exactly 1,075 unique validation cells to the 1,075 spatially-resolved training cells. CellTRIP imputed spatial coordinates match the shape of the observed training data, overcoming the main challenge of imputing spatial coordinates directly (Figure 3a). When evaluating cell type distributions by cortical layer, we observe that CellTRIP has comparable performance to all other methods and mimics cell type enrichments (Section S2.2) from the reference dataset closely (Figure 3b, Supplementary Figure S6). Among the surveyed methods, CellTRIP layer classifications have the lowest matrix MSE (Section S2.2) for excitatory cells (CellTRIP 0.042, CellTREK 0.066, Tangram 0.108, Cytospace 0.073, Supplementary Figure S7). CellTRIP outperforms other methods when classifying L5 PT and L6b cell types as layers 5 and 6b, respectively. In particular, CellTRIP achieves the highest AUROC on single-layer classifiers for layers L5 and L6b (Supplementary Figure S8). Layer L6b has traditionally been difficult to identify and analyze given its relatively thin width, heterogeneous cell distributions, and lack of agreed-upon marker genes^35^. We define a layer score for each reference and imputed single-cell (Section S2.2) and observed that CellTRIP has significantly different (p < 10^*−*4^, Mann-Whitney U test) distributions between most excitatory cell types, as well as astrocytes and L2/3, which was only identified by CellTRIP and Tangram (Figure 3c).

**Figure 3.**
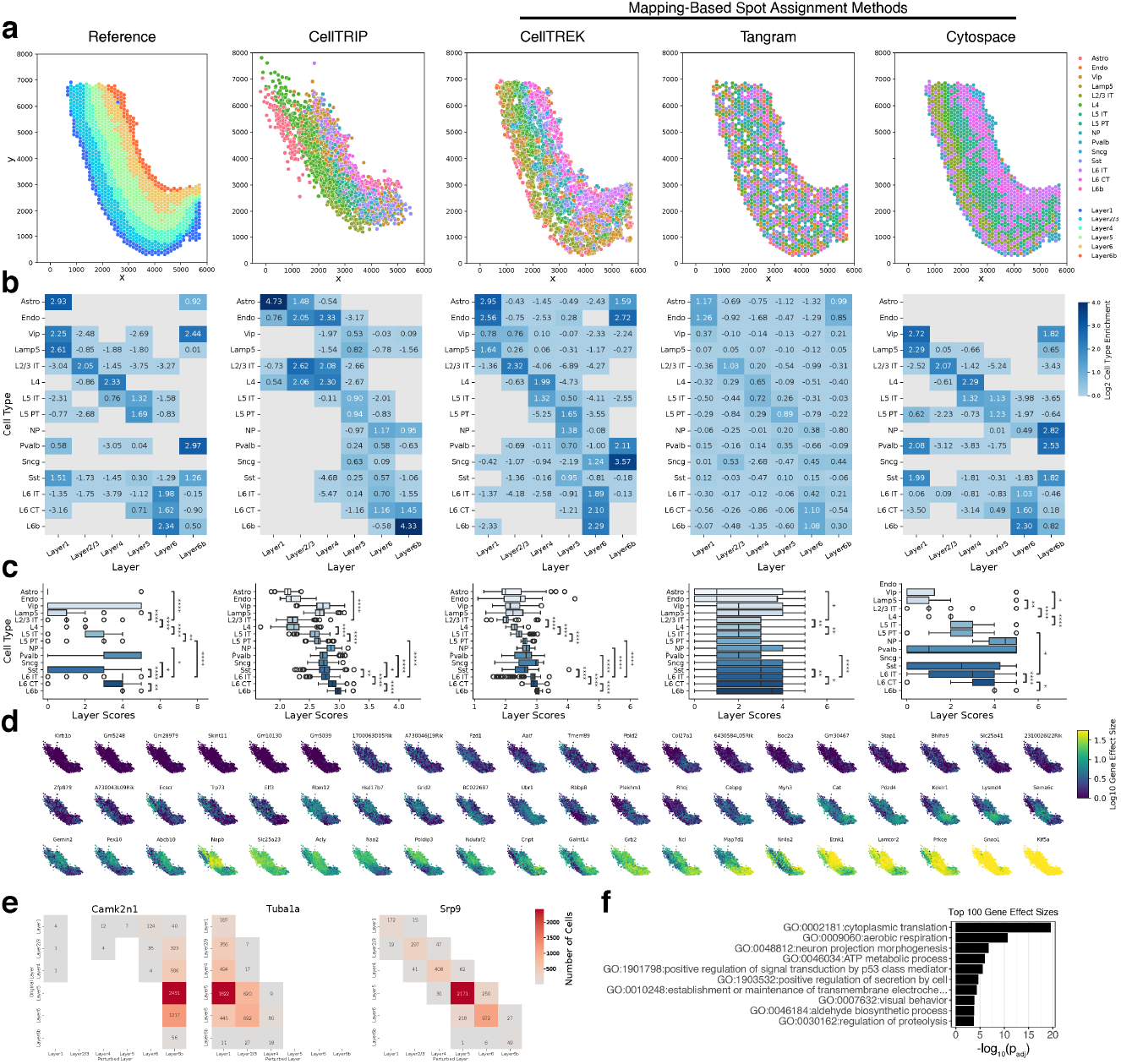
Single-cell spatial imputation and gene prioritization in the adult mouse frontal cortex. For computational details, see Section S2.2. **a.** Reference spatial transcriptomics training data ^29^ from Wei et al. ^30^ and imputed spatial coordinates of single cells from independent mouse cortical scRNA-seq data^25^. We compared CellTRIP with mapping-based spot assignment methods including CellTREK^30^, Tangram^31^, and Cytospace^32^. **b**. Cell type enrichment (log^2^ transformed) of inferred reference subclasses (left) compared with imputed coordinates from (a) across cortical layers. **c**. Single-cell layer score distributions per cell type for methods shown in (a) to assess separability of subclasses by cortical layer, particularly for excitatory neurons. Significances of distribution differences between inferred adjacent cortical layers are computed using a one-tailed Mann-Whitney U test and annotated as *: p < 5 ×10^*−*2^, **: p < 1 × 10^*−*2^, ***: p < 1 × 10^*−*3^, ****: p < 1 × 10^*−*4^. **d**. Imputed spatial coordinates from CellTRIP, colored by gene effect sizes after short (.5s) in-silico gene knockdown by CellTRIP. 60 random genes were chosen for visualization. Plots were sorted using hierarchical clustering of genes by single-cell effect sizes. **e**. Number of cells transitioning to (x-axis) and from (y-axis) cortical layers under gene knockdown by CellTRIP for the two top genes with the highest number of layer transitions, along with one representative gene for average layer transitions. Knockdown was fully simulated for the top 1,000 genes by effect size. **f**. Gene set enrichment of top 100 genes by number of transitioning cells.

We then used CellTRIP for in-silico perturbation prediction of gene knockdown and calculated spatial gene effect sizes (Section S2.2) on validation data across all genes (Figure 3d). The distribution of gene effect sizes is heavily right skewed, indicating that many genes have little to no effect on the spatial imputation (Supplementary Figure S9). We then checked cell movement across layers under knockdown for the top 1,000 genes by effect size (Section S2.2). We examine the layer transitions of two such genes with high numbers, *Camk2n1* and *Tub1a*, as well as one with an average number of transitioning cells, *Srp9* (Figure 3e). Under CellTRIP-predicted perturbation, both *Camk2n1* and *Tub1a* cause catastrophic layer migration towards L6b and L1, respectively. Knockdown of *Camk2n1* impairs maintenance of longterm memory by preventing typical post-retrieval autophosphorylation^36^. *Tub1a* also plays a critical, noncompensated role in neuronal saltatory migration. Performing functional enrichment on the top 100 genes by gene effect size, we obtain the significant term neuron projection morphogenesis, which is heavily interrelated with neuronal migration (Figure 3f). interestingly, we see an additional enriched term for long-term memory, which aligns with the prioritization of *Camk2n1*.

### 4.3 Predicting Developmental Gene Expression Changes After Drug Treatment in Cancer Cells

Next, we sought to evaluate the accuracy CellTRIP perturbation trajectories, particularly for intermediate timepoints. To do so, we used a dataset consisting of 13,713 cells, 5,992 of which have been treated with trametinib and subsequently measured for gene expression at 3, 6, 12, 24, and 48 hour timepoints. During training, we held out the 24 hour timepoint completely for both control and trametinib.

CellTRIP demonstrated superior perturbation prediction from dimethyl sulfoxide (DMSO) control to trametinib at 48 hours for all cell lines. Before benchmarking, we verified that the cell line distributions were similar across observed DMSO and trametinib cells (Supplementary Figure S10a). For CellTRIP and GEARS, we simulated drug perturbation by simulating knockdown on the two main targets of trametinib, *MAP2K1* and *MAP2K2*, whose expressions appeared to decrease monotonically over the course of treatment (Supplementary Figure S10b). We utilized three postprocessing adjustment strategies when evaluating CellTRIP, namely *No Adjustment, Steady-State Adjustment*, and *PCA Adjustment* (Section S2.3). We see that PCA-adjusted CellTRIP (MSE = 3.87, Pearson delta = 0.650) outperforms CPA^37^ (MSE = 5.52, Pearson delta = −0.092) and GEARS^38^ (MSE = 68.74, Pearson delta = 0.414) in both MSE and Pearson delta (Figure 4a). We note that the comparison methods only predict one perturbation at a time (e.g., *control →Tram_48hr*) while CellTRIP can evaluate multiple timepoints in one perturbation trajectory.

**Figure 4.**
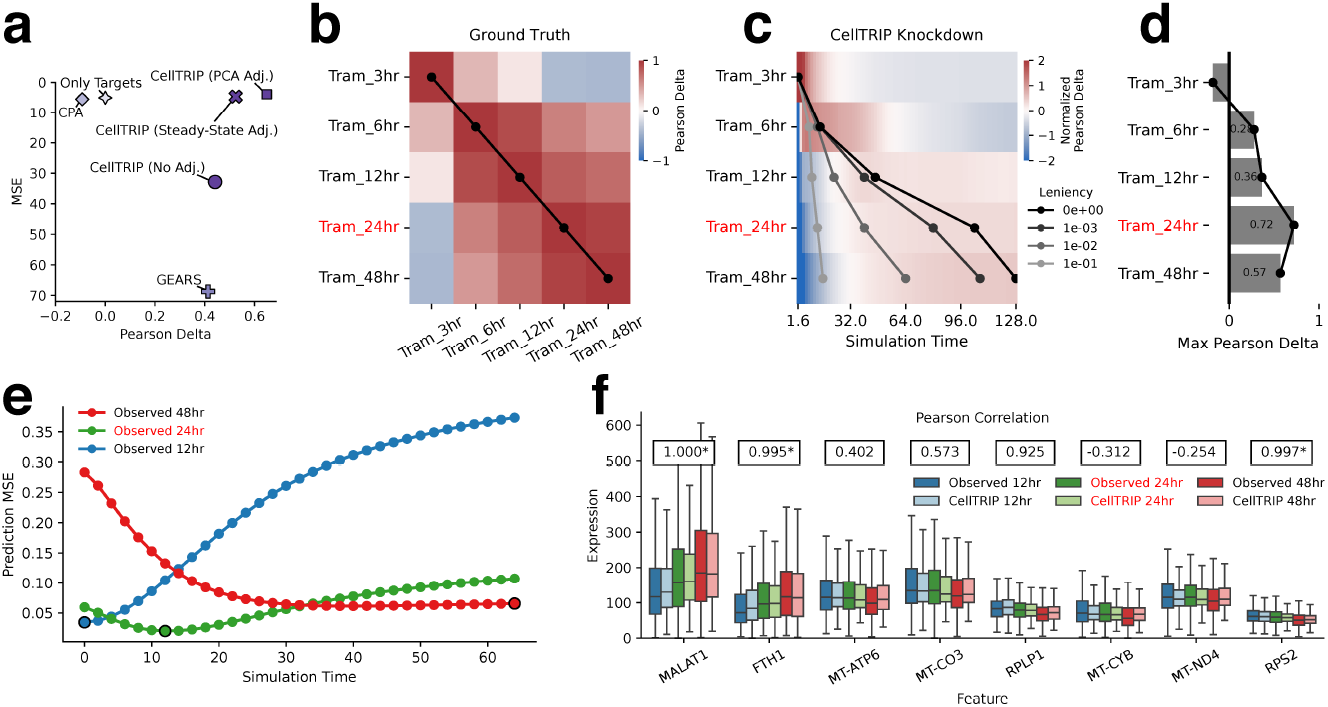
Gene knockdown perturbation prediction and missing timepoint recovery for trametinib in cancer cells^7^. **a.** CellTRIP knockdown prediction performance against comparable methods using Pearson delta (See Section S2.3). **b**. Pearson delta between recorded timepoints after trametinib perturbation, using the unperturbed control (DMSO) at 48 hours as a baseline. **c**. Normalized Pearson delta between timepoints from (e) and CellTRIP perturbation after knockdown of *MAP2K1* and *MAP2K2* genes to the same values as trametinib at 48 hours. Lines indicate the earliest points where the value of a sliding mean window with size four reaches within *leniency* of the maximal mean Pearson delta for each row. **d**. Maximal Pearson delta of CellTRIP-predicted knockdown for each timepoint of trametinib perturbation. **e**. Wasserstein distance (EMD) between observed 12, 24, and 48 hour timepoints after trametinib perturbation and CellTRIP transition states between 12 and 48 hours. Observed data was limited to the top 512 principal components. Expected closest timesteps are outlined for each series, as well as a manually chosen timestep for 24 hours to be used in (d). **f**. Observed and predicted gene expression for 12, 24, and 48 hour timepoints.

Using the continuous and interactive environment from CellTRIP, we can predict perturbation trajectories over long periods of time, and thereby predict multiple timepoints simultaneously. To do so, we first ran the environment to steady state on control cells. Then, we knocked down *MAP2K1* and *MAP2K2*. The knockdown perturbation provided a trajectory from 0 to 48 hours, which we then compared to our expectations. Specifically, we compute the Pearson delta between different timepoints of trametinib perturbation to determine the similarity between perturbed states, and, correspondingly, the expected distribution of these timepoints in our simulated trajectory (Figure 4b). We note that the latter four timepoints (6 hours onward) are more similar to each other than the 3 hour timepoint. Additionally, closer timepoints are generally more similar. Performing the same analysis with Pearson delta between perturbation timepoints and CellTRIP timesteps, we see a similar trend to that of the ground truth (Figure 4c). Importantly, peaks of the computed Pearson deltas appear in each perturbation timepoint sequentially, suggesting that the CellTRIP trajectory is nearing each perturbation timepoint in the correct order. CellTRIP is also able to achieve a high Pearson delta of 0.72 at 24 hours, sharing positive results with all but the 3 hour timepoint (Figure 4d).

To showcase the continuous perturbation trajectory from CellTRIP, we imputed trametinib-perturbed gene expression at 24 hours, using 12 and 48 hour measurements as referenceo. Following the methodology in Supplementary Figure S1, we generated 948 pseudocells for the 12 and 48 hour timepoints, 948 being the minimum cell count between the two, and estimated pseudocell correspondence between the timepoints using discrete optimal transport. We then ran the environment to steady state on the 12 hour pseudocells before replacing cell expression with the corresponding 48 hour cells, giving us a trajectory between the two timepoints. We observed that the distribution of pseudocells began closest to that of the 12 hour timepoint, approached the held out 24 hour timepoint at the midpoint of the simulation, then finally converged to the 48 hour timepoint, as expected (Figure 4e). We manually picked a timestep from this trajectory to represent the 24 hour timepoint. We then compared CellTRIP-predicted and actual gene expression at the beginning, manually-chosen, and ending timepoints (Figure 4f). Meadian CellTRIP expressions across all timepoints were significantly correlated for three of the top eight genes by change from 12 to 48 hours, and positively correlated with three others.

### 4.4 Spatiotemporal Reconstruction of *Drosophila* Embryonic Development

We further trained a single CellTRIP model on a single-cell spatiotemporal gene expression dataset in *Drosophila* larvae, demonstrating its capability for imputation in both temporal and spatial domains, as well as for perturbation prediction. This dataset measures both gene expression and the spatial coordinates of 155,684 single cells across five developmental stages (two embryonic and three larval). Note that we held out the intermediate developmental stage *E16-18h_a* from training for imputation evaluation.

We applied CellTRIP to the task of imputing spatial coordinates from measured gene expression (Section 3.4.1). CellTRIP spatial imputation outperformed MLP- and KNN-based methods (Section S2.4). In methods other than CellTRIP, we observed overfitting to later larval stages, evidenced by the stretched distribution of cell spatial coordinates in embryonic and early larval stages (Figure 5a). CellTRIP spatial imputation performance was comparable to other methods on most training stages, but had the lowest prediction MSE (CellTRIP 90.41, MLP 333.86, KNN 360.33) and highest classification accuracy of 10 tissue types (CellTRIP 0.237, MLP 0.231, KNN 0.207, Baseline 0.100) on the heldout developmental stage *E16-18h_a* and in a majority of constituent tissue types (Figure 5b and Supplementary Figure S11).

**Figure 5.**
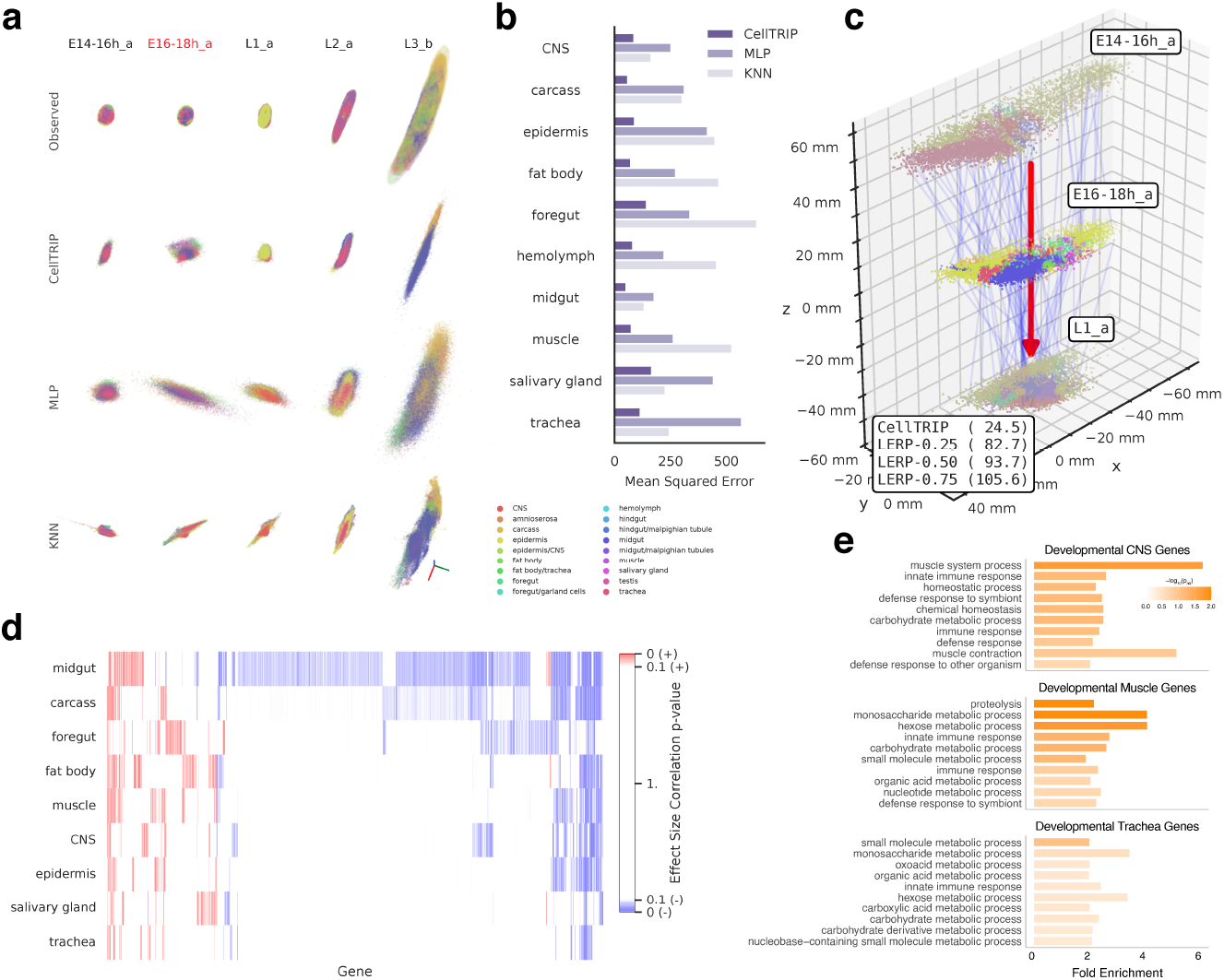
Spatiotemporal developmental stage imputation, recovery, and knockdown in developing *Drosophila*. **a.** Visualization of observed and predicted spatial coordinates for fly embryonic development^8^ across developmental timesteps (columns) and methods (rows), colored by cell annotations. Stage *E16-18h_a* (red) was not included in training. **b**. Mean squared error for select annotations compared across methods. **c**. Heldout developmental stage recovery using CellTRIP. Wasserstein distance (EMD) is computed between predicted and observed cell distributions, including baseline linear interpolation (LERP) at a few transitional fractions. Further methodological details can be found in Section 3.4.1 and Figure S1. **d**. Heatmap of genes (x-axis) with developmentally-correlating perturbation significance values as predicted by CellTRIP, segmented by annotation (y-axis). Stronger Pearson correlation significance is indicated by higher opacity while positive and negative correlation is indicated by red and blue coloration, respectively. **e**. Gene set enrichment of top 200 CellTRIP developmental genes from select cell annotations.

We then performed temporal imputation to infer the spatial trajectory of cells between known developmental timepoints, and subsequently recover the cellular spatial distribution of our held out intermediate developmental stage (Section 3.4.2, Figure S1). First, we use optimal transport to match pseudocells between the stages immediately preceding and following *E16-18h_a*, namely, *E14-16h_a* and *L1_a*. We then used CellTRIP to compute a spatial trajectory between the two timepoints, selecting an intermediate trajectory timestep to recover the cell spatial distribution of *E16-18h_a* (Figure 5c). CellTRIP outperformed linear interpolation (LERP) using the Wasserstein distance (EMD) between observed and imputed spatial coordinates (CellTRIP 24.5, LERP 82.7, Section S2.4).

We further utilized CellTRIP to make in-silico gene perturbation predictions to prioritize tissue type developmental genes. In particular, we calculated gene effect sizes (Section 3.4.3) for the top 2,000 highly-variable genes by tissue type and developmental stage. We then computed Pearson correlations of our gene effect sizes with developmental stages to prioritize developmental genes (Section S2.4, Figure 5d).

We focused on tissue types where developmental progression is morphologically well-characterized and mechanistically linked to stage remodeling. Based on this criterion, we selected central nervous system (CNS), muscle, and tracheal tissue types, representing neuronal maturation ^39^, skeletal muscle development^40^, and respiratory tube branching ^41^, respectively (Figure 5e). In CNS, the top 10% developmental genes by correlation were significantly enriched for innate immune response (p < 2.36 × 10^*−*4^), a key aspect of the *Drosophila* CNS which plays a key role in preventing harmful effects from injury, infection, and neurodegenerative diseases^42^. Performing a similar analysis in muscle tissue, we obtain a top enrichment of proteolysis (p < 2.12 × 10^*−*5^), p_adj_ < 1.07 × 10^*−*2^)). Muscle proteolysis is critical for maintaining glucose supply under starvation conditions and may share pathways responsible for muscle atrophy^43^. Lastly, many metabolic processes are prioritized for tracheal tissue (small molecule metabolic process, p < 1.05 × 10^*−*4^), and metabolic depression under starvation uses insulin as to stimulate tracheal progenitor cells for regeneration or remodeling ^44^.

## 5 Discussion

We have shown the outperformance and predictive capabilities of CellTRIP for trajectory interpolation, imputation, and perturbation prediction on a diverse range of temporal and spatial datasets. CellTRIP is also interactive, biologically interpretable, and can provide importance estimates with respect to individual genes, spatial locations, and developmental stages. To our knowledge, this is the first application of reinforcement learning to generate virtual cell environments based on single-cell omics data. We also contribute to existing reinforcement literature by extending PopArt^21^ to support input layer normalization and weight rescaling. CellTRIP has the distinct advantage of being interactive. CellTRIP can be generalized to other dynamic datasets, such as disease progression or tissue-level analysis with bulk expression data.

CellTRIP has some limitations. In Section 4.2, we note that imputation of the uncorrected validation data yields spatial coordinate predictions shifted from the observed data. We choose to apply batch correction to the validation dataset. Further analysis would be needed, however, to determine how much of this shift was caused by batch effects or was driven by true spatial expression variance. We note that stage *L3_b* from Section 4.4 displays a tendency of CellTRIP to distribute cells only across the primary spatial axis, likely due to the focus of CellTRIP on generality across time points. If sufficient information exists in gene expression alone, then this may be solved by running more simulations in parallel during training, allowing for a more representative training set each backward pass. We run CellTRIP with a low-dimensional environment space because the training time for a continuous reinforcement learning model scales rapidly. This is a well-known side effect of continuous output in PPO models and can be solved with either greater memory size or discretization of the model outputs. We found that the environment space performs better when diagonal action vectors remain unnormalized. This has the consequence that cells in the environment can move faster diagonally, which is unideal. The runtime of CellTRIP also typically scales with the number of simulated and visible cells, as expected. However, distributed computing for forwards and backwards passes and user-adjustable inference settings mitigate this runtime limitation significantly. CellTRIP needs to employ additional strategies to ensure minimal usage of both memory and VRAM throughout the running process. CellTRIP can also be extended to predict cell death, proliferation, or the spread of disease in tissue samples through modifications to the environment space and policy actions.

## Notes

### Competing Interest Statement

The authors have declared no competing interest.

### Funding

- NSF: Career 2144475
- NIH: RF1MH128695

