## Supplemental Materials for "Inferring virtual cell environments using multi-agent reinforcement learning"

### S1 Model Training

CellTRIP training is an iterative process consisting of the aforementioned simulation and backpropagation. To train a CellTRIP model for a given single-cell dataset, new environment spaces are randomly initialized in cell position and velocity, then simulated to completion. The cell states of the environment, as well as the actions and corresponding rewards for each simulation timestep, are recorded per-cell for later use in backpropagation. These computations are referred to as a memory for each cell, so for each timestep, $n_c$ memories will be stored. Each completed simulation is referred to as an episode. On average, around $300$ episodes are simulated to train each CellTRIP model for our applications in this study. The simulations can also be filtered such that each environment space trains only cells from a particular cell group (e.g., specific developmental stage or disease state).

After a user-defined number of memories are generated, by default $1{,}000{,}000$, the model performs a policy update (Supplementary Section S1.1). The update itself is split into $80$ epochs, by default, within each of which the model iterates over all stored memories. Each epoch is divided into batches which update policy weights using a subset of the data in each epoch, typically $10{,}000$ memories each. For the applications in this paper, we use the models after $800$ policy updates. To lower memory and computational requirements during training, CellTRIP employs additional strategies such as agent vision limitation, batch computation, and Monte Carlo sampling, which are user-adjustable (Supplementary Section S8.3). Additionally, several memory storage optimizations are utilized by default (Supplementary Section S8.1).

#### S1.1 Policy PPO, Entropy, and Critic Losses

The CellTRIP policy considers three main losses to optimize cell action prediction: Proximal policy optimization (PPO) loss[1], entropy loss, and critic loss. The PPO loss reinforces cell actions which lead to higher environmental rewards, the entropy loss encourages exploration of the environment space, and the critic loss improves the ability of the policy to predict future environmental rewards. We denote the policy weights $\theta$, with a corresponding actor network $\pi_\theta$, which predicts cell actions $\pi_\theta(\mathcal{S}^t)$. An additional critic network $\omega_\theta$, is trained concurrently but separately to the actor network with nearly identical structure, predicting the per-cell reward for each state, $\omega_\theta(\mathcal{S}^t)$. The actor and critic networks share weights for all but the decider module, which is replaced with a simple fully connected layer with a single-feature output for the critic. The full loss function is notated as below

$$\ell_{\text{policy}} = \ell_{\text{ppo}} + 10^{-3}\,\ell_{\text{entropy}} + \ell_{\text{critic}}. \tag{S1}$$

The default loss coefficients are shown, but can be adjusted by users.

To calculate the PPO loss, we first define a maximal change parameter $\epsilon$, defaulted to $0.2$. We iterate the policy using Supplementary Equation S2 on cell actions $\Delta V^t$ and states $\mathcal{S}^t$ for each timestep

$t$.

$$\ell_{\text{ppo}} = \min\left(\frac{\pi_\theta(\Delta V^t|\mathcal{S}^t)}{\pi_{\theta_{\text{old}}}(\Delta V^t|\mathcal{S}^t)}\hat{a}^t,\ \text{clip}\left(\frac{\pi_\theta(\Delta V^t|\mathcal{S}^t)}{\pi_{\theta_{\text{old}}}(\Delta V^t|\mathcal{S}^t)}, 1-\epsilon, 1+\epsilon\right)\hat{a}^t\right). \tag{S2}$$

where $\pi_\theta(\Delta V^t|\mathcal{S}^t)$ denotes the probability of policy $\pi_\theta$ taking action $\Delta V^t$ given state $\mathcal{S}^t$ and $\pi_{\theta_{\text{old}}}$ is the policy at the beginning of backpropagation. $\hat{a}^t$ notates the advantage, which is typically the difference between the actual discounted environmental reward, $\vec{d}^t$, and expected discounted reward, $\omega_{\theta_{\text{old}}}(\mathcal{S}^t)$, as predicted by the critic network before backpropagation. In CellTRIP, however, we use generalized advantage estimation (GAE)[2] to compute the advantage directly (Supplementary Section S1.2).

The actor network has an additional loss to incentivize raising $\sigma$, thereby encouraging policy exploration of the environment space. This loss is inversely proportional to the entropy from the sampling method in Section 3.3,

$$\ell_{entropy} = \frac{n_d}{2}\log(2\pi e\sigma^2). \tag{S3}$$

We choose the weight $10^{-3}$ for the $\ell_{entropy}$ term based on empirical tuning, targeting convergence of $\sigma$ to $0$ after $400$ policy updates.

The loss function for the critic network is the Smooth L1 Loss from PyTorch[3]. This loss behaves similarly to MSE for smaller errors, but more linearly as the error increases. Given the inferred discounted rewards $\tilde{d}^t = \hat{a}^t + \omega_{\theta_{\text{old}}}(\mathcal{S}^t)$, the loss takes on the following form:

$$\ell_{\text{critic}} = \begin{cases} .5(\tilde{d}^t - \omega_\theta(\mathcal{S}^t))^2 & \text{if } |\tilde{d}^t - \omega_\theta(\mathcal{S}^t)| < 1, \\ |\tilde{d}^t - \omega_\theta(\mathcal{S}^t)| - 0.5 & \text{otherwise,} \end{cases} \tag{S4}$$

where $\omega_\theta(\mathcal{S}^t)$ is the predicted discounted reward from the updated critic network.

All losses are aggregated according to Supplementary Equation S1. $\ell_{\text{policy}}$ is then used to optimize $\theta$, the policy parameters, using typical gradient descent. CellTRIP has an additional option to limit Kullback-Leibler divergence of actions from $\pi_\theta$ to $\pi_{\theta_{\text{old}}}$, but this is not utilized in the base model. We refer to the optimized weights as $\hat{\theta}$, and the trained actor and critic as $\pi_{\hat{\theta}}$ and $\omega_{\hat{\theta}}$, respectively.

#### S1.2 Generalized Advantage Estimation and Reward Propagation

We use generalized advantage estimation, or GAE[2], to reduce variance in reward during policy training at the expense of additional bias. This is performed through estimation of $\hat{a}^t$ rather than predicting $\vec{d}^t$ directly. Allowing temporary losses to realize future rewards is a key component of successful reinforcement learning approaches. Without GAE, we might compute the reward of a state using the sum of the reward and discounted future rewards.

$$\vec{d}^t = \vec{r}^t + \gamma\vec{d}^{t+1} = \vec{r}^t + \gamma\vec{r}^{t+1} + \gamma^2\vec{r}^{t+2} + \dots, \tag{S5}$$

where $\gamma$ is a user-defined discount rate for future rewards, defaulting to $0.99$. Rewards are propagated per cell, per episode.

However, $\vec{d}^t$ is highly variant with respect to future actions, making cause-and-effect associations harder to pinpoint. To reduce this variance in favor of bias, we utilize GAE. Rather than computing the discounted reward directly, we instead compute the advantage. GAE is defined as the weighted average of a $k$-step advantage estimator $\hat{a}^{(k),t}$,

$$\vec{\zeta}^t = \vec{r}^t + \gamma\omega_\theta(\mathcal{S}^{t+1}) - \omega_\theta(\mathcal{S}^t), \tag{S6}$$

$$\hat{a}^{(k),t} = \sum_{l=0}^{k-1}\gamma^l\vec{\zeta}^{t+l} = \vec{r}^t + \gamma\vec{r}^{t+1} + \dots + \gamma^{k-1}\vec{r}^{t+k-1} + \gamma^k\omega_\theta(\mathcal{S}^{t+k}). \tag{S7}$$

Note that Supplementary Equation S7 approaches Supplementary Equation S5 as $k \to \infty$. The exponentially-weighted average of these $k$-step advantage estimators is the following,

$$\hat{a}^t = (1-\lambda)(\hat{a}^{(1),t} + \lambda\hat{a}^{(2),t} + \lambda^2\hat{a}^{(3),t} + \dots) = \sum_{l=0}^{\infty}(\gamma\lambda)^l \vec{\zeta}^{t+l}, \tag{S8}$$

where $\lambda$ is a user-defined discount rate for higher-order $k$-step estimators, defaulting to $0.95$.

In practice, Supplementary Equation S8 is computed until the end of each simulation. Note that $r^t$ is $0$ for $t > T$, where $T$ is the last simulation timestep. This means that both $\vec{d}^t$ and $\hat{a}^{(k),t}$ will generally decrease as $t$ increases where $t + k - 1 > T$. To remedy this, we employ a novel form of bootstrapping, which computes $\omega_\theta(\mathcal{S}^{T+1})$ for use in computing $\vec{\zeta}^T$. In other words, we use only use the $1$-step advantage estimator, $\hat{a}^{(1),T}$ to compute $\hat{a}^T$. Although this introduces bias, our rewards tend to decrease over time, so the effect is minimized. These $\hat{a}^t$ estimations are utilized in Supplementary Equation S2 to produce the optimal actor, $\pi_{\hat{\theta}}$. The advantage is also used to calculate $\tilde{d}^t$, which is then used to train the optimal critic, $\omega_{\hat{\theta}}$, as well.

#### S1.3 Adaptive Input and Output Rescaling

Reinforcement learning, and particularly PPO, is generally sensitive to changes in input and reward distributions. Intuitively, rewards from the environment and the critic model will increase with training as the actor improves. In certain applications, the distribution of inputs will change as well. For CellTRIP, we observed that mean absolute distance from the origin, and thereby the overall magnitude of the environment space coordinates for each cell, decrease as training continues and the model becomes more refined. This also results in shrinkage of the distribution of cells in the environment space. These distributional changes constitute additional challenges for model convergence, so much so that CellTRIP models would often experience performance collapse or a lack of convergence without intervention.

PopArt[4] proposes a solution to the reward distribution shift. First, while performing a policy update, we estimate the first and second moments, $\vec{\mu}^t$ and $\vec{\nu}^t$, of the reward distribution online,

$$\vec{\mu}^t = (1-\beta)\vec{\mu}^{t-1} + \beta\,\vec{\eta}^{t,1}, \quad \vec{\nu}^t = (1-\beta)\vec{\nu}^{t-1} + \beta\,\vec{\eta}^{t,2}, \tag{S9}$$

where $\vec{\eta}^{t,1}$ and $\vec{\eta}^{t,2}$ are the mean and mean of squares of the critic network estimations, $\omega_\theta(\mathcal{S}^t)$, across a batch and $\beta$ is a tunable weight parameter, $3 \times 10^{-3}$ by default. The standard deviation can then be estimated with $\vec{\sigma}^t = \sqrt{\vec{\nu}^t - (\vec{\mu}^t)^2}$. We allow the critic network to compute a normalized output, and replace $\tilde{d}^t$ in Supplementary Equation S4 with

$$d^t_{\text{norm}} = \frac{\tilde{d}^t - \vec{\mu}^t}{\vec{\sigma}^t}. \tag{S10}$$

This is referred to as adaptively rescaling targets (Art). Note that we can also unnormalize critic outputs using $\vec{\sigma}^t\omega_\theta(\mathcal{S}^t) + \vec{\mu}^t$.

However, this technique shifts $\ell_{\text{critic}}$, making reward or advantage estimation more difficult. So, we add a component for preserving outputs precisely (Pop). Specifically, using our estimated mean and variance, for each update of $\mu^t$ and $\nu^t$, we also update the weights and bias of the final critic layer, $W_{\text{output}} \in \mathbb{R}^{n_{\text{hidden}} \times n_{\text{output}}}$ and $\vec{b}_{\text{output}} \in \mathbb{R}^{n_{\text{output}}}$,

$$W^t_{\text{output}} = \frac{\vec{\sigma}^{t-1}}{\vec{\sigma}^t} \odot W^{t-1}_{\text{output}}, \quad \vec{b}^t_{\text{output}} = \frac{\vec{\sigma}^{t-1}\vec{b}^{t-1}_{\text{output}} + \vec{\mu}^{t-1} - \vec{\mu}^t}{\vec{\sigma}^t}, \tag{S11}$$

where division occurs element-wise and $\odot$ indicates elementwise multiplication where vectors are broadcast over the first dimension. Note that this update cancels out that from Supplementary Equation S10.

In other words, for the same input, the updated model will produce the same output. In practice, we apply PopArt to the output layers of our pinning modules (Section 3.4.1) in addition to the output layer of the critic, which requires adapting PopArt for multiple features, as we did above.

We extend the concepts from CellTRIP to input layers, specifically those of the actor/critic and pinning modules. To do so, we need only replace Supplementary Equation S11 with

$$(W^t_{\text{input}})^T = \frac{\vec{\sigma}^t}{\vec{\sigma}^{t-1}} \odot (W^{t-1}_{\text{input}})^T, \quad \vec{b}^t_{\text{input}} = \vec{b}^{t-1}_{\text{input}} + \frac{\mu^t - \mu^{t-1}}{\sigma^{t-1}} (W^{t-1}_{\text{input}})^T \tag{S12}$$

where $W_{\text{input}} \in \mathbb{R}^{n_{\text{input}} \times n_{\text{hidden}}}$ and $\vec{b}_{\text{input}} \in \mathbb{R}^{n_{\text{hidden}}}$ are the weights and biases of the input layer, respectively. We arrived at this formulation by first inverting the change to $W^t_{\text{output}}$ in Supplementary Equation S11, then solving for $\frac{\vec{x}-\vec{\mu}^{t-1}}{\sigma^{t-1}} W^{t-1}_{\text{input}} + \vec{b}^{t-1}_{\text{input}} = \frac{\vec{x}-\vec{\mu}^{t}}{\sigma^{t}} W^{t}_{\text{input}} + \vec{b}^{t}_{\text{input}}$.

### S2 Datasets, Evaluation, and Analysis

#### S2.1 Simulated Cell Development and Knockdown

We train a CellTRIP model on $1{,}500$ single cells generated using Dyngen[5], withholding $20\%$ of the data as validation. The data consists of seven cell states labeled `A` through `G` which develop according to a predetermined lineage, illustrated by Figure 2b. This lineage contains two major bifurcations, one at cell state `C`, which allows for differentiation to terminal cell state `G` or another bifurcation at cell state `D`, which can differentiate to terminal cell states `E` or `F`. Cells are annotated based on their trajectory in the format `sX{mid/end}_sY{mid/end}`, where `X` and `Y` are cell states and `{mid/end}` indicate bifurcations and terminal cell types, respectively. Each cell state has anywhere between $5$ and $14$ modules, where a single module contains up to nine transcription factors (TFs). In total, there are $4{,}800$ cell features split evenly across gene and protein expression, making two modalities with $2{,}400$ features each. Before training with CellTRIP, we apply log-normalization to the gene expression modality.

To assess the quality of the CellTRIP-reconstructed modalities, we compare to several methods with similar dimensionality reduction strategies. Namely, PCA, autoencoders, JAMIE[6], and scScope[7]. PCA, autoencoder, and scScope methods were run only on gene expression data, while JAMIE was trained on both gene expression and protein counts. For performance, we rely on two metrics, pairwise mean square error (Pairwise MSE) and label transfer accuracy (LTA). The former is the MSE between the inter-cell distance matrices for the simulated and reconstructed gene expression. The latter is equivalent to the accuracy of a k-nearest neighbors classifier (n=$10$) trained on labels from the simulated gene expression when applied to the reconstructions from each method.

To perform module knockdown, the CellTRIP environment is first simulated to steady state on the full simulation dataset. Then, we simultaneously knock down the expression of all transcription factors within the module to zero. The resultant trajectories can then be visualized or quantified by gene effect size, as in Section 3.4.3. We also simulate knockdown of individual TFs, taking the mean across gene modules to assess predicted versus expected gene effect sizes across trajectories. Note that we compute gene effect sizes in the environment space, rather than expression space.

#### S2.2 Spatial Transcriptomics in the Adult Mouse Cortex

All models are trained on $1{,}075$ spatially-resolved cells from the adult mouse across six cortical layers[8], subset to the frontal cortex[9]. For validation, we impute $4{,}785$ cells from an adult mouse cortical cell taxonomy, annotated by cell type[10]. Both datasets are adjusted to have per-cell gene counts of $10{,}000$ before being log normalized. Before being processed by CellTRIP, the datasets are further standardized and reduced to $512$ principal components. We apply additional batch correction to the validation dataset, matching the means of each gene with those in the training data.

We use all comparison spot assignment methods to assign the validation expression to spots from the training data, and thereby associate cell types from the validation expression with cortical layers from the training spatial coordinates. For CellTRIP, we instead use a KNN regression model with $200$ neighbors, trained to infer ordinally-arranged cortical layers (i.e., L1 is $0$, L2/3 is $1$, etc.) from the training spatial coordinates, and compute the median and IQR for each layer. We then infer cortical layers from the CellTRIP predicted validation spatial coordinates and take the nearest layer median, after dividing by the IQR, as the predicted layer. We perform this procedure to attempt to remove bias against CellTRIP as a regression model when evaluating against spot assignment methods. After association, we calculate enrichment scores by dividing the fraction of each cell type within a layer by the overall fraction of cells belonging to said cell type. Before visualization, we apply a $\log_2$ transform to these enrichments.

To evaluate model predictions, we computed the MSE between the row-normalized matrix of layer associations and ground truth across all excitatory cell types. For our ground truth matrix, we assumed each excitatory cell type belonged only to its associated layer (e.g., L5 IT belongs to L5). We used a similar layer association strategy when determining per-cell-type and per-layer accuracies. To include inhibitory cell types, we generated a cell type reference dataset by transferring cell type labels from the validation to the training data using only gene expression with Scanpy[11]. We also used an smFISH reference dataset consisting of $2{,}360$ cells from the primary visual cortex[12,13] to establish ground truth cell type distributions per cortical layer.

We additionally computed layer scores for each validation cell per method by computing the mean ordinally-arranged cortical layer of the nearest $1{,}000$ cells, weighted by the inverse of their Euclidean distances from the cell. We used a one-tailed Mann-Whitney U test to determine significant differences between cell type layer score distributions in adjacent layers, empirically determined based on layer score ordering across all methods. This empirical ordering was also used for ordering of the cell type enrichment heatmap subclasses.

After computing per-cell gene effect sizes for single-gene knockdown, we visualized randomly-sampled perturbation results. These results are sorted by hierarchical clustering from SciPy[14], using per-cell gene effect sizes for the feature dimension. Using these gene effect sizes, we prioritized $1{,}000$ genes for extended simulation (from $1$s to $128$s). With these extended simulations, we further counted cells transitioning across layers under knockdown to identify spatially-relevant genes. When performing gene set enrichment on any extracted gene lists, we refer to the FDR-adjusted (Benjamini-Hochberg) $p$-value as $p_{\text{adj}}$.

#### S2.3 Trametinib Treatment of Cancer Cell Lines

The dataset from McFarland et al.[15] contains gene expression for $13$ drugs at $3$, $6$, $12$, $24$, and $48$ hours after treatment, as well as the untreated cells at $48$ hours. We construct a training set from the untreated cells, dimethyl sulfoxide (DMSO), and trametinib. DMSO is a common organic solvent used in medical applications. Trametinib itself is typically administered as a DMSO solvate, and targets *MEK1* and *MEK2* enzymes. Before training the CellTRIP model, we apply sample count ($10{,}000$ samples) and log-normalization.

We begin by benchmarking CellTRIP against existing perturbation methods CPA[16] and GEARS[17]. In particular, we simulate knockdown of *MAP2K1* and *MAP2K2* genes, which encode the enzymes *MEK1* and *MEK2*. In CellTRIP, this implies running the DMSO $48$ hour cells to steady state, then replacing tex expressions of *MAP2K1* and *MAP2K2* with their means in the trametinib treated cells at $48$ hours. We apply three post-processing strategies to compute the CellTRIP perturbation. *No Adjustment* utilizes the difference between the final imputed state from the simulation and the original untreated data as the predicted perturbation delta. *Steady-State Adjustment* takes the difference between the imputed final and beginning states of the knockdown simulation to get the perturbation delta. *PCA Adjustment* uses the difference between the imputed final state and the PCA-reconstructed untreated expression, limited to 512 principal components. To evaluate the perturbations from each method, we use MSE and

Pearson delta, which measures the correlation between gene expression changes under predicted and true perturbations.

We then evaluate the CellTRIP perturbation trajectory from DMSO to trametinib at $48$ hours. We obtain a ground-truth heatmap by computing the Pearson delta between each trametinib perturbation timepoint, annotated by a line contacting the maximal Pearson delta in each row. To perform evaluation for CellTRIP, we compute Pearson delta between all perturbation timepoints (y-axis) and the CellTRIP-predicted timesteps (x-axis). For visualization, we row-normalize the heatmap. Ideally, the ordinality of peaks in each row will match that of the ground truth. We annotate this plot with lines for different *leniency* values. Specifically, we compute mean Pearson deltas for each row with a sliding window of $4$ timesteps, spanning roughly $12.8$ seconds of simulation time. Each line contacts the first point in each row which is within *leniency* of the maximal value. For this analysis, we utilize another CellTRIP model trained on raw expression counts rather than log-normalized counts.

Further, we follow the interpolation procedure described in Supplementary Figure S1 to obtain a trajectory between $12$ and $48$ hour expressions. Specifically, we generate pseudocells equal to the minimal number of cells in either timepoint, $948$, and establish correspondence using optimal transport. We then simulate the $12$ hour cells to steady state and replace their expression profiles with those of the $48$ hour pseudocells, giving us a trajectory which can be evaluated for distance from the observed $12$, $24$, and $48$ hour timepoints. To evaluate the trajectory, we compute MSE at each simulation timestep. To evaluate CellTRIP perturbation in individual genes, we take the genes with the $8$ greatest median differences in expression between $12$ and $48$ hour timepoints, plotting their expression for $12$, $24$, and $48$ hour timepoints alongside CellTRIP predictions. We additionally compute pearson correlation between the observed and CellTRIP-predicted median expressions for each timepoint.

#### S2.4 Spatial Transcriptomics for *Drosophila* Development

The data from Wang et al.[18] consists of $155{,}684$ single-cells spanning across five developmental stages, two embryonic and three larval, of *Drosophila*. The dataset contains $11{,}703$ common genes across developmental stages with accompanying spatial coordinates. Before being input to CellTRIP, both the gene expression data and spatial coordinates were normalized. The gene expression data was then reduced to $512$ principal components using PCA. To evaluate spatial imputation (Section 3.4.1) from gene expression, we generally used MSE and tissue type classification accuracy as performance measures. Tissue type classification accuracy being the accuracy with which a KNN classifier trained on the spatial coordinates of the observed developmental stages can classify tissue types of imputed single-cells. We compare to an MLP with layers $11{,}703 \times 100 \times 3$ and ReLU activation and a KNN regression model with $10$ neighbors using Scikit-Learn[19]. To compare performance of our spatiotemporal imputation (Section 3.4.1 and Section 3.4.2), we use linear interpolation, which is a weighted mean between spatial coordinates of our observed initial and terminal states. We perform this estimation at several *transition fractions* $(0.25, 0.5, 0.75)$ which serve as the weights for the initial stage spatial coordinates. To evaluate the interpolated stages, we use the Wasserstein distance (EMD)[20]. Briefly, the Wasserstein distance documents the cost, or total movement, of an optimal transport plan. For visualization, we classify tissue types of the interpolated stage using a KNN classifier with $10$ neighbors trained on the original intermediate stage.

In computing the developmental gene prioritizations, we first measure the spatial gene effect size (Section 3.4.3) for all developmental stages. The knockdown simulation is performed on all cells simultaneously, but recorded for each annotated region separately. This allows for per-region gene significance estimations. After normalizing all samples across cell annotation and developmental stage, we can compute Pearson correlations revealing prioritized genes with monotonic developmentally-variant gene effect sizes.

### S3 Spatial Imputation

To determine our matrix $R$ and vector $\vec{\tau}$, we begin by computing centroids,

$$\overrightarrow{\text{cent}}_{\rho k} = \frac{1}{N} \sum_{i=1}^{n_c} \rho^{(k)}(\vec{x}_{e,i}), \quad \overrightarrow{\text{cent}}_{Mk} = \frac{1}{N} \sum_{i=1}^{n_c} \vec{m}_i^{(k)}. \tag{S13}$$

where $\overrightarrow{\text{cent}}_{\rho k}$ and $\overrightarrow{\text{cent}}_{Mk}$ are row vectors. We then use singular value decomposition on the covariance matrix, which is a product of the centered matrices, to determine the optimal rotation.

$$U \Sigma V^T = (\rho^{(k)}(X_e) - \overrightarrow{\text{cent}}_{\rho k})^T (M^{(k)} - \overrightarrow{\text{cent}}_{Mk}). \tag{S14}$$

After solving for $\vec{\tau}$ in $R\rho^{(k)}(X_e) + \vec{\tau} = M^{(k)}$, we finally have

$$R = UV^T, \quad \vec{\tau} = \overrightarrow{\text{cent}}_{Mk} - \overrightarrow{\text{cent}}_{\rho k} R. \tag{S15}$$

This technique allows CellTRIP to impute both modalities that have individual cell significance and those that are only significant relative to other cells. Theoretically, this can introduce noise in the training process, as changes to individual cell predictions can affect the accuracy of others. However, in practice, we observed no such degradation.

### S4 Important Notes for Training Models

- If a model runs out of memory during backpropogation, it is usually a result of the `minibatch _memories` parameter being too high.
- If one reward dominates between pinning and velocity, they can be adjusted using the environment keyword arguments `reward_pinning` and `penalty_velocity`.
- Although it may lead to more stable training, using the flag `dont_sync_across_nodes` greatly decreases computation time, due to the large overhead from uploading memories to the Ray object store.

### S5 Methodology Clarifications

#### S5.1 Modality Source and Targets

We split modalities $k \in K$ across $K_{\text{sources}}$ and $K_{\text{targets}}$ according to the application. As an example, integration applications will have $K_{\text{sources}} = K_{\text{targets}} = K$, indicating that all modalities are input and imputed by the policy and pinning modules. In the case of single-modal imputation, $|K_{\text{targets}}| = 1$ and $K_{\text{targets}} \vee K_{\text{targets}} = K$ where $\vee$ denotes the exclusive union.

#### S5.2 Pinning Module Training and Synchronization

Pinning modules train largely according to the procedure in Section 3.4.1. There are a couple important implementation details which we will document here. Primarily, by default, the pinning modules only train on memories from simulation times past $32$ seconds. This time is chosen as the maximal time for convergence, after which the cells should not make any large movements in the environment space. Without this restriction, the pinning module trains on earlier simulation states, which may be volatile and biologically uninformative. Secondarily, this restriction makes distributed synchronization more difficult.

Specifically, in a given batch, it is possible that one learner (See Supplementary Section S6) will update its pinning modules while another does not. To prevent the update process from hanging and keep the pinning modules synchronized across learners, we allow the non-updating pinning module to perform an update with $0$ gradient. To prevent diminishing the gradient from only one pinning module, we additionally synchronize a tensor between learners to determine the effective world size (i.e., how many pinning modules have data to use in an update) to correctly determine the mean gradient update from all actively updating pinning modules. Pinning modules are trained after the actor and critic for a default of five epochs with $1{,}024$ epoch size and $64$ batch size.

### S6 Distributed Processing

The distributed forward of CellTRIP is implemented using Ray[21]. In particular, we use an autoscaling AWS cluster consisting of one `m5.large` instance which deploys `g5.2xlarge` instances with NVIDIA A10G GPUs (22Gb VRAM) and 32Gb of RAM as needed.

To start training, a manager node, typically with no GPU, coordinates processes and writes checkpoints and logs to s3 or the local machine using the `RecordBuffer` class. The manager deploys any number of worker agents across any number of machines. Typically, each worker machine will have a GPU. One worker on each machine is assigned to be the head worker, which is responsible for synchronizing model weights across machines using the NVIDIA Collective Communication Library (NCCL). Depending on user preference, workers are then assigned roles as learners and/or runners. The former is responsible for backpropagation and is assigned only to the head node of each machine, while the latter is only responsible for simulation. By default, we assign two workers to be both learners and runners.

Upon initialization, the head workers average model weights across machines and propagate the synchronized weights to any other workers on the same machine. Note that model weights include statistics and optimizers, specifically, parameters for input and output normalization, as well as an update counter. All workers simulate independently until a user-defined number of memories have been generated. By default, this number is $1{,}000{,}000$ divided by the number of workers. Importantly, we stagger the lengths of each simulation randomly (See Section 3.1) so that memories are not over-centralized on a single simulation segment during backpropagation.

At this point, if the user did not pass the `dont_sync_across_nodes` flag, memories are synchronized across all nodes. Otherwise, memories are only synchronized to the learner worker on each machine. Backpropagation then begins on all learner nodes with scaled epoch and batch sizes according to the number of learners. Each weight update, the learner nodes synchronize gradients. Optionally, users can choose to only synchronize in certain intervals. However, in this case, all weights need to be synchronized rather than just gradients. In the case of the pinning module, synchronization is slightly more complicated (See Supplementary Section S5.2).

Users can also decide intervals of numbers of updates, at which the manager node will save a checkpoint. CellTRIP is also able to resume training from checkpoints across any Ray cluster configuration.

### S7 Optional CellTRIP Functionality

#### S7.1 Knockdown Modules

In addition to simply replacing features with a target value as in Section 3.4.3, which we refer to as *feature clamping*, we include additional capabilities to perform perturbation and gene knockdown. Namely, *inverse feature clamping* can be used when PCA is utilized for preprocessing. The technique calculates the closest PCA-reduced representation by MSE using least-squares for which the representation, after inverting PCA, recovers the target feature at the target value. In other words, instead of knocking a feature down and applying PCA (like *feature clamping*) this technique will ensure that the recovered feature from the PCA-reduced representation is knocked down. Lastly, *move towards targets* is a more invasive approach that applies a force to each action from the model, directed towards the area in the environment space corresponding to the target feature value. This is often used in tandem with *inverse feature clamping*.

In practice, we use *inverse feature clamping* and *move towards targets* for all perturbation applications aside from Figure 4a, where we use traditional *feature clamping*. This is generally a decision that should be made based on manual analysis of the model performance.

##### S7.2 Training Stages

In the case of particularly high-dimensional environment spaces or complex data, users may wish to train in stages, gradually introducing each loss from Equation 2 over the course of training. This is achievable through user-configured hyperparameters. If enabled, the model progresses through stages using an early stopping criterion. By default, if the mean reward over the past $3$ model updates has not increased in $6$ updates, the stage advances. Although this technique is not utilized due to the stabilility of CellTRIP on our analyzed datasets, it may prove useful for more complex applications.

##### S7.3 Discrete Representation

CellTRIP has the additional capability to be run with a multi discrete action space, outputting $-1$, $0$, or $1$ for each environment dimension. The environment then limits the resulting action vector to a user-defined magnitude, typically $1$.

#### S8 Runtime Optimizations

##### S8.1 Memory Storage Optimizations and Sampling

Several complications arise when implementing the model, many of which have to do with memory and efficient computation. Chief among these is storing memories, which is integral to the generality of the model after training.

In the simplest approach, the total number of memories stored between updates scales with both the number of timesteps between updates and the number of cells being simulated in the environment. Each cell has a different state for the same timestep, as the cells are processed differently depending on whether they are the main or neighboring cells. Each state can be represented as a matrix of size $n_c \times (2n_d + \sum_k n_{mk})$.

As a conservative example, suppose we run the model forward for $5,000$ timesteps with $1,000$ cells and $2$ modalities with $3,000$ features each (and a $16$-dimensional latent space). We then have $40$M states totaling ~$109$TB (for 32-bit data). In code, this is all stored in our *memory handler object*.

However, much of this data is redundant as the modal representations of cells rarely change over the course of training. Therefore, we can cut the modal features from the stored states. These modal features in the state matrix are replaced with indices corresponding to a vector containing the original modal data, greatly reducing redundancy. When a memory is accessed, the memory handler object appends the modal data back onto the state, which allows for normal processing by the model. This alone cuts the memory requirement to ~$317$GB in the example above. Using better indexing techniques, we can optimize further.

There is still redundancy in the state storage. Namely, the same information is stored for a cell $n_c$ times over, as the inputs to the model are different in each case. We can instead store the state only once, then format the input to the model based on which memory is accessed. When a memory is indexed, the memory handler object scans through the stored states to determine to which cell the index corresponds. Then, the corresponding cell is separated from the state matrix and the two are returned after appending their modal data. In the example above, this further reduces memory usage to only ~$324$MB. Altogether, we have reduced memory usage by a factor of ~$35$k.

It is worth noting that this strategy comes at the cost of indexing time. For this reason, backward passes have strategies implemented to minimize the need to pull from the memory handler object. More details about these strategies are found in Supplementary Section S8.4.

Additionally, CellTRIP has the capability to run on datasets of any size, and includes methods to pull from online sources, including s3, when sampling new data. This flexibility comes with increased latency and memory requirements. As such, for computing preprocessing classes, we limit CellTRIP to $10{,}000$ samples by default.

#### S8.2 Batch Tensor Construction

As input, the CellTRIP model takes three tensors. In order, these tensors are the cell tensor (cells $\times$ features), neighbor tensor (cells $\times$ features), and the attention mask (cells $\times$ cells). The attention mask is a boolean tensor documenting masked attentions for each cell in the cell tensor. During the model backward, these are batched into three 3D tensors. In the case of jagged memories, or memories with unequal numbers of cells, batching memories into a tensor requires padding to the maximal number of cells across all sampled memories. This padding can incur a high memory cost depending on the training environment, especially in the case of batching few samples from many different timepoints.

#### S8.3 Batch Computation

In instances where modalities contain many features or the number of cells is relatively large, it may not be feasible to compute all cell actions at once depending on the amount of VRAM available. Due to the independent action computation of each node, the model forward may be split by simply computing only a few cells at a time. Although this increases the time of the forward significantly, the process is far more memory efficient.

In cases with many cells, it can also become necessary to only sample a subset of neighbor cells in addition to the target cell. This can be achieved by simply omitting the neighbor cells from the computation described in Section 3.3. The approach does have an effect on the 'vision' of the agent, as its view of the state is limited to the chosen neighbor cells. However, this necessitates additional storage in the memory object. Namely, the cells which are taken for computation must be recorded. This complicates the indexing approach discussed in Supplementary Section S8.1. In the default case, CellTRIP uses the hash of the current state matrix to seed a random function to decide the nodes which are visible to the agent.

#### S8.4 Backward Sampling

During the model backward, there are several strategies which can be used to balance memory usage with efficient computation. We have implemented a flexible and adaptive system which attempts to maximize throughput given user-defined limitations on the processing. Within this system, we first define a *pool* (default size unlimited) of memories. This is the group of memories which is used across all epochs during the model backward. We then choose memories for each epoch (default size $100{,}000$) from the memories within the pool. We then split the epoch into batches (default size $10{,}000$). If a batch is too large to be processed all at once, minibatches (default limited to $1{,}000{,}000$ memories) are further subset from the batches and run individually. The gradients of all minibatches are then added before iterating the model at the end of each batch. This is generally referred to as gradient accumulation.

### S9 Supplementary Tables

| Symbol | Definition |
|---|---|
| $n_c$ | Number of cells |
| $n_d$ | Environment space dimensionality |
| $M^{(k)}$ | Single-cell data for modality $k$ |
| $\vec{m}_i^{(k)}$ | Single-cell data for modality $k$ and cell $i$ |
| $t$ | Simulation timesteps |
| $\Delta t$ | Simulation step time |
| $K, K_{\text{sources}}, K_{\text{targets}}$ | Lists of all, source, and target modalities |
| $X^t$ | Cell position in environment space |
| $\vec{x}_i^t$ | Cell position in environment space and cell $i$ |
| $V^t$ | Cell velocity in environment space |
| $\vec{v}_i^t$ | Cell velocity in environment space and cell $i$ |
| $\Delta V^t$ | Cell action in environment space |
| $\Delta \vec{v}_i^t$ | Cell action in environment space and cell $i$ |
| $\mathcal{S}^t$ | Cell states, consisting of positions, velocities, and source modalities |
| $\vec{r}^t$ | Total environmental reward |
| $\vec{r}^t_{\text{pinning}}$ | Pinning reward |
| $\vec{\delta}^t$ | Stepwise pinning reward |
| $\delta_i^t$ | Stepwise per-cell pinning reward |
| $\rho^{(k)}$ | Pinning module for modality $k$ |
| $\vec{r}^t_{\text{velocity}}$ | Velocity penalty |
| $\vec{r}^t_{\text{action}}$ | Action penalty |
| $\vec{a}_i^t$ | Cell embeddings |
| $E_a$ | Cell encoder |
| $\vec{b}_i^t$ | Neighbor embeddings |
| $E_b$ | Neighbor encoder |
| $S^t$ | Cell summary embeddings |
| $\vec{s}_i^t$ | Cell summary embedding for Cell $i$ |
| $E_s$ | State encoder |
| $\sigma$ | Policy action standard deviation |
| $\pi_\theta$ | Policy actor with weights $\theta$ |
| $\omega_\theta$ | Policy critic with weights $\theta$ |
| $\theta_{\text{old}}$ | Weights before policy update |
| $\epsilon$ | PPO maximal change parameter |
| $\hat{a}^t$ | Advantage |
| $\vec{d}^t$ | Discounted reward |
| $\hat{d}^t$ | Predicted discounted reward |
| $\hat{d}^t_{\text{old}}$ | Predicted discounted reward from critic before policy update |
| $\tilde{d}^t$ | Inferred discounted reward |
| $\ell_{\text{ppo}}$ | PPO loss |
| $\ell_{\text{entropy}}$ | Entropy loss |
| $\ell_{\text{critic}}$ | Critic loss |
| $\gamma$ | Reward discount factor |
| $\hat{a}^{(k),t}$ | $k$-step advantage estimator |
| $\vec{\zeta}^t$ | Intermediate reward estimation |
| $\lambda$ | $k$-step estimator discount factor |

| Symbol | Definition (Continued) |
|---|---|
| $T$ | Terminal simulation timestep |
| $\beta$ | New batch weight for online reward distribution estimation |
| $\mu^t$ | Online estimation of reward distribution first moment |
| $\nu^t$ | Online estimation of reward distribution second moment |
| $\eta^{t,1}$ | Mean of batch reward estimations |
| $\eta^{t,2}$ | Mean of square batch reward estimations |
| $n_{\text{input}}$ | Size of input layer |
| $n_{\text{hidden}}$ | Size of hidden layer |
| $n_{\text{output}}$ | Size of output layer |
| $W_{\text{input}}$ | Weights of input layer |
| $W_{\text{output}}$ | Weights of output layer |
| $\vec{b}_{\text{input}}$ | Bias of input layer |
| $\vec{b}_{\text{output}}$ | Bias of output layer |
| $C$ | Euclidean distance matrix between initial and terminal stages for interpolation |
| $Q$ | Transition matrix calculated via optimal transport |
| $\phi$ | Perturbation function |
| $\phi_g$ | Knockdown function on feature $g$ |
| $\ell_{\text{pinning},k}$ | Modality $k$ pinning module loss |
| $R$ | Rotation matrix |
| $\vec{t}$ | Translation vector |
| $\overrightarrow{\text{cent}}_{\rho k}$ | Centroid for pinned cell positions |
| $\overrightarrow{\text{cent}}_{Mk}$ | Centroid for observed cell positions |
| $U, \Sigma, V$ | Components of singular value decomposition |

Table S1: Definitions of commonly-used mathematical symbols.

### S10 Supplementary Figures

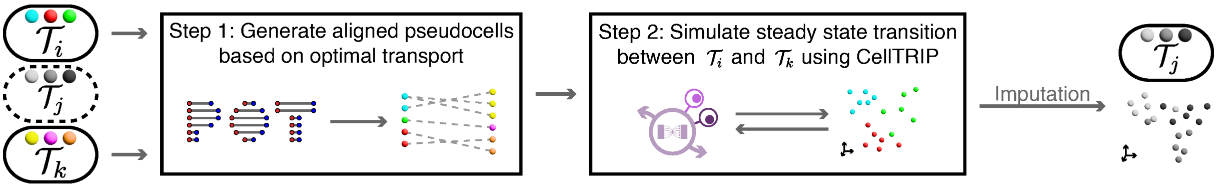


Figure S1: Inference of intermediate timepoints using CellTRIP. To infer intermediate developmental stages (e.g. timepoints, cell types) with unmatched cells, CellTRIP uses optimal transport[22] to compute pseudocells between the observed initial and terminal states. The initial environment is simulated to steady state. Then, the initial environment input modalities are replaced with the terminal modalities and simulated to steady state once more. The resultant transition states contain the CellTRIP predicted developmental progression from the initial to terminal stages.

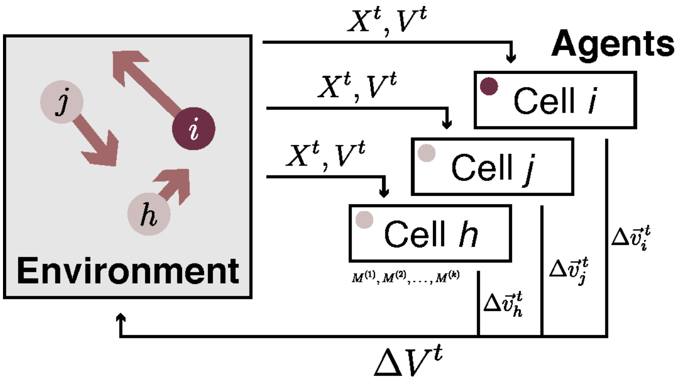


Figure S2: Illustration of the environment update loop. Cells, which are represented by agents in the environment space, consider the positions, velocities, and modalities of all other cells before determining an action. All actions are sent to the environment simultaneously, moving to the next simulation timestep.

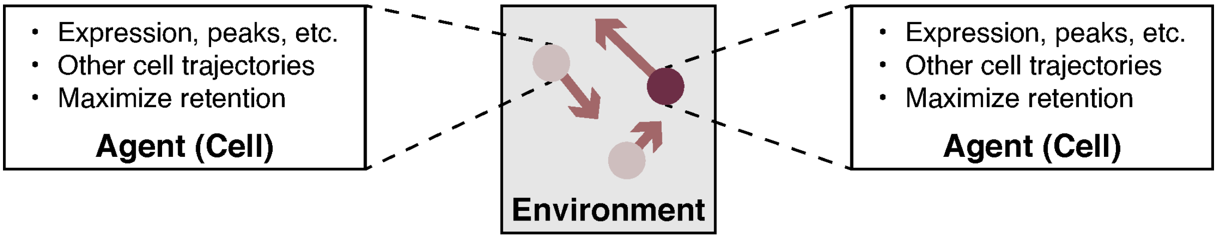


Figure S3: Illustration of the multi-agent reinforcement learning environment. Cells are represented as independent agents controlled using a shared policy, which seeks to maximize rewards for each agent.

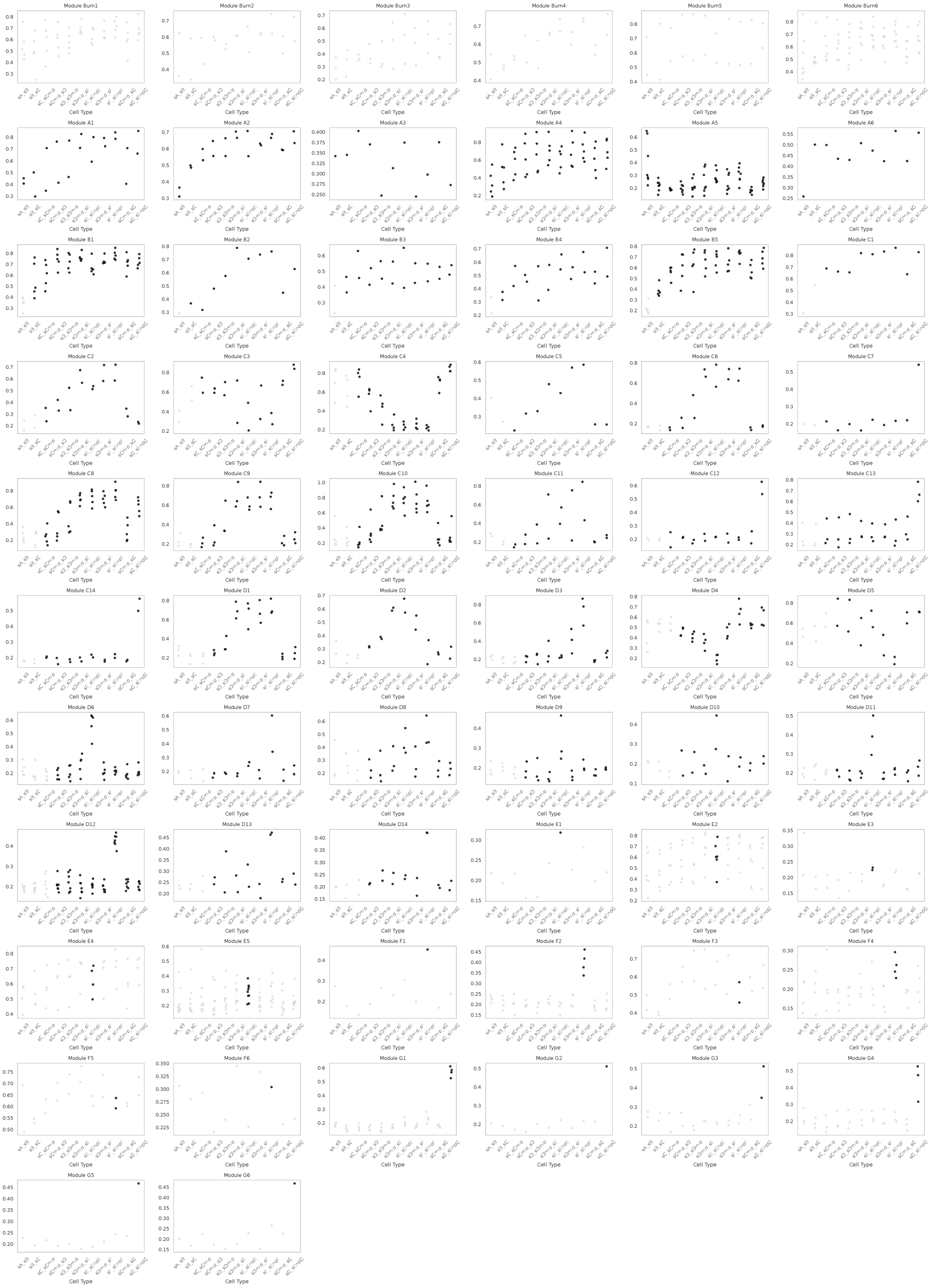


Figure S4: CellTRIP-predicted gene effect sizes for knockdowns on transcription factors (TFs) from all modules in Dyngen-generated[5] simulation data. TF gene effect sizes for downstream trajectories (child cell states) are black, while others are gray.

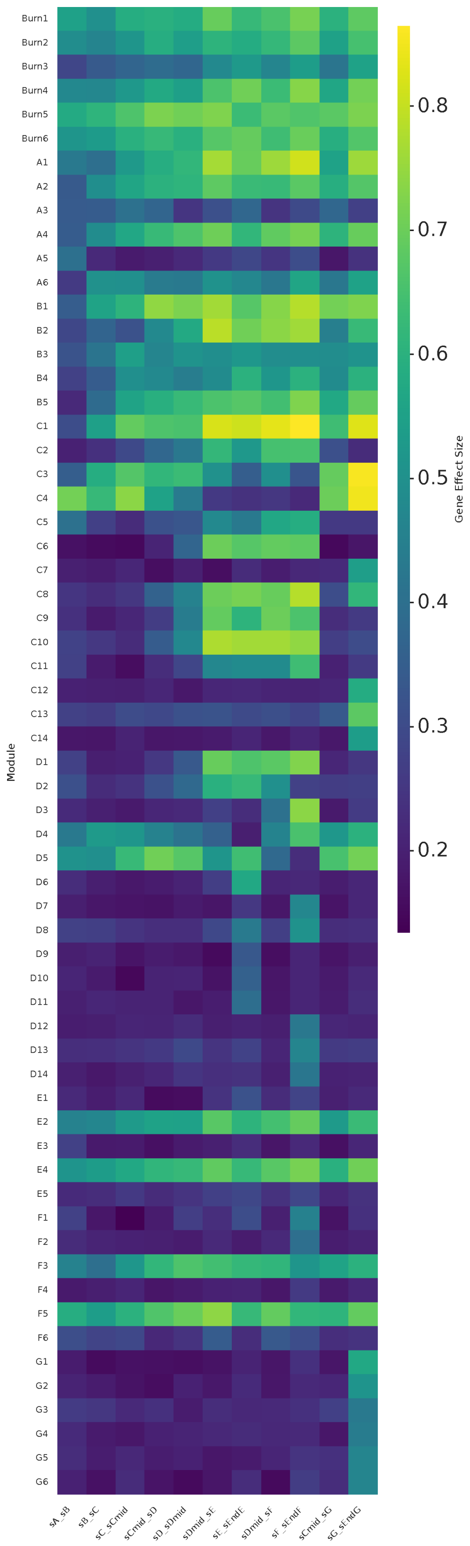


Figure S5: Heatmap of mean CellTRIP-predicted gene effect sizes in environment space by cell trajectory (x-axis) and all modules (y-axis). Extended heatmap from Figure 2f.

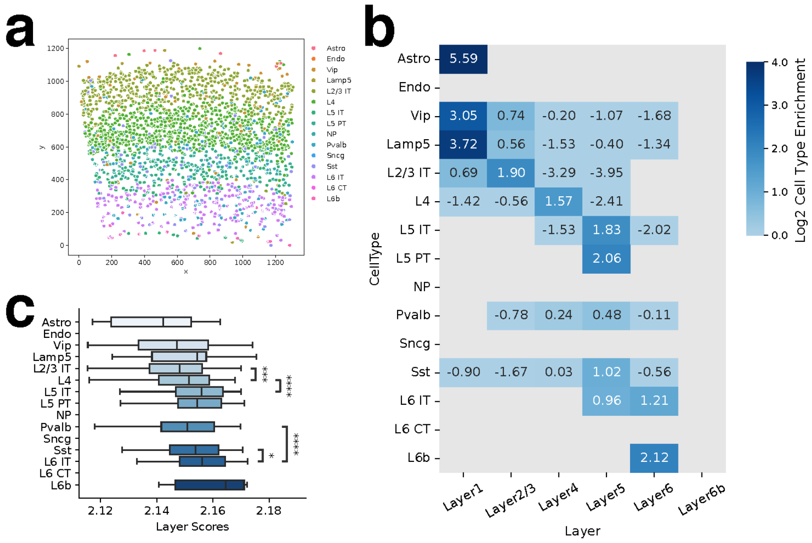


Figure S6: Reference smFISH data[12,13] with 2,360 cells from the mouse primary visual cortex. **a.** Spatial coordinates, annotated by cell type. **b.** Cell type enrichment ($\log_2$ transformed) of each cell type across cortical layers. **c.** Single-cell layer score distributions per cell type. Significances of distribution differences between inferred adjacent cortical layers are computed using a one-tailed Mann-Whitney U test and annotated as *: $p < 5 \times 10^{-2}$, **: $p < 1 \times 10^{-2}$, ***: $p < 1 \times 10^{-3}$, ****: $p < 1 \times 10^{-4}$.

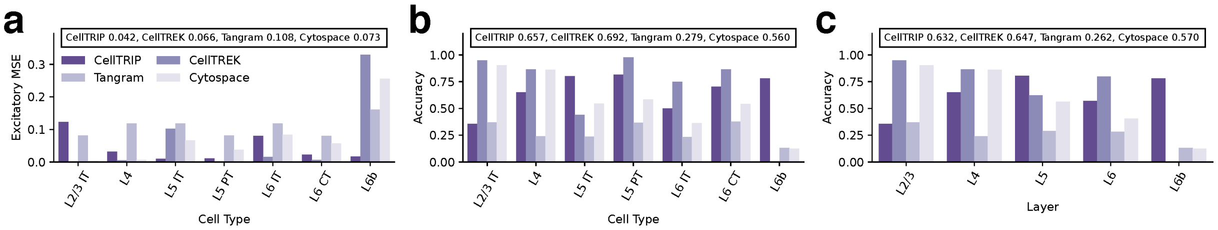


Figure S7: Performance comparison between CellTRIP, CellTREK[9], Tangram[23], and Cytospace[24] on validation gene expression data from the adult mouse frontal cortex[10]. **a.** MSE of row-normalized layer predictions from each method versus ground truth by cell type (See Supplementary Section S2.2). **b.** Accuracy of layer predictions from each method by cell type. **c.** Accuracy of layer predictions from each method by cortical layer.

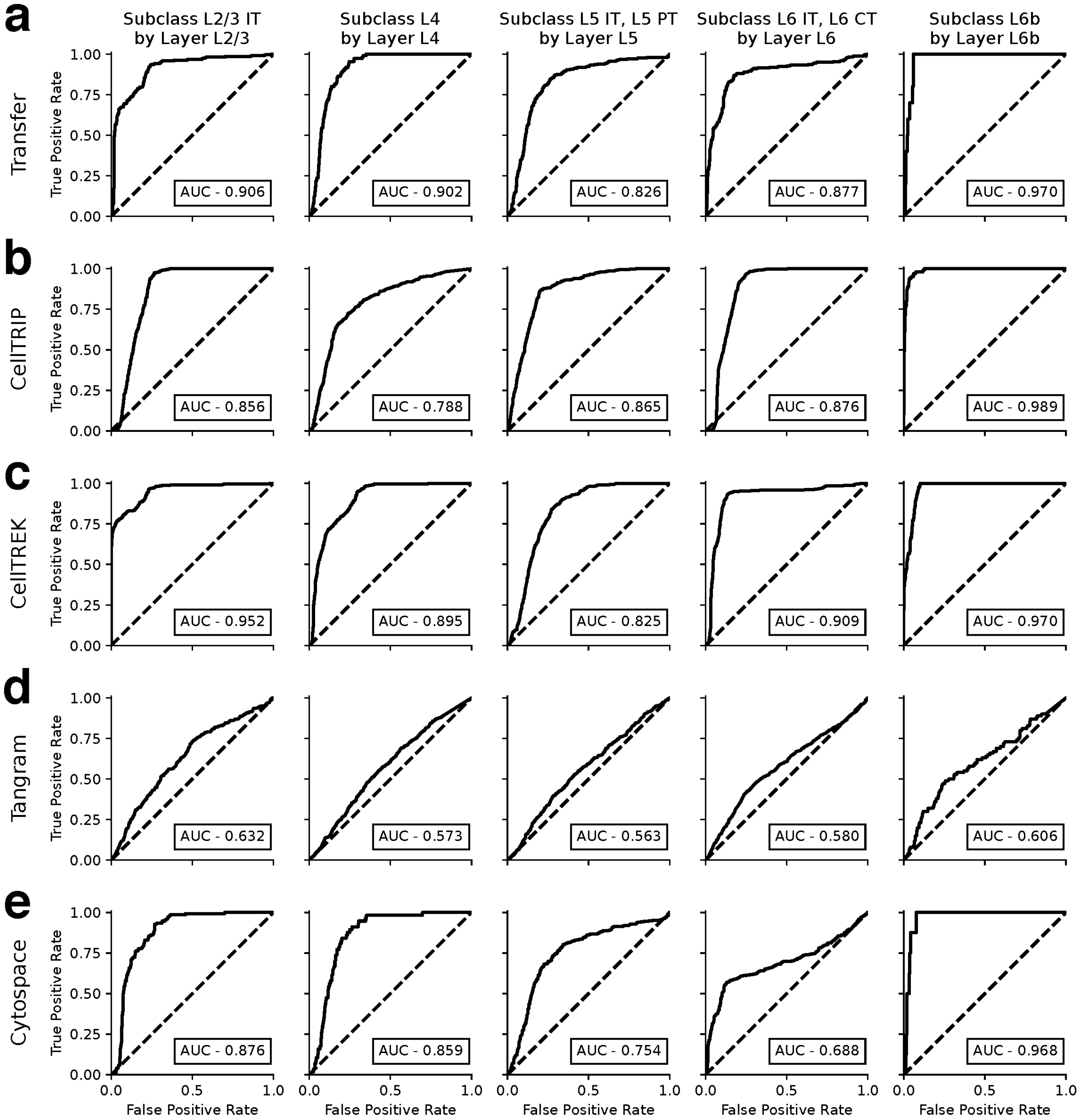


Figure S8: Classifier ROC curves on excitatory cell types from the adult mouse frontal cortex[10] for reference label transfer data (a), CellTRIP (b), CellTREK[9] (c), Tangram[23] (d), and Cytospace[24] (e).

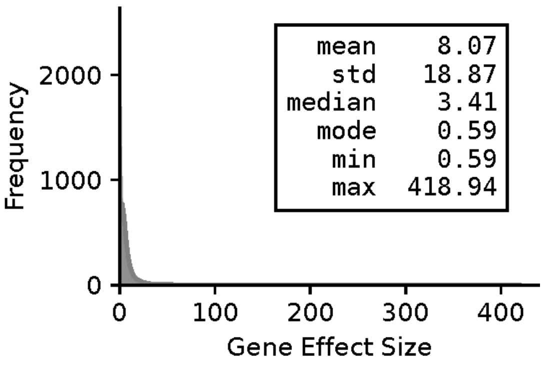


Figure S9: Histogram of mean gene effect sizes after single-gene knockdown on validation gene expression data from the adult mouse frontal cortex[10] for all genes.

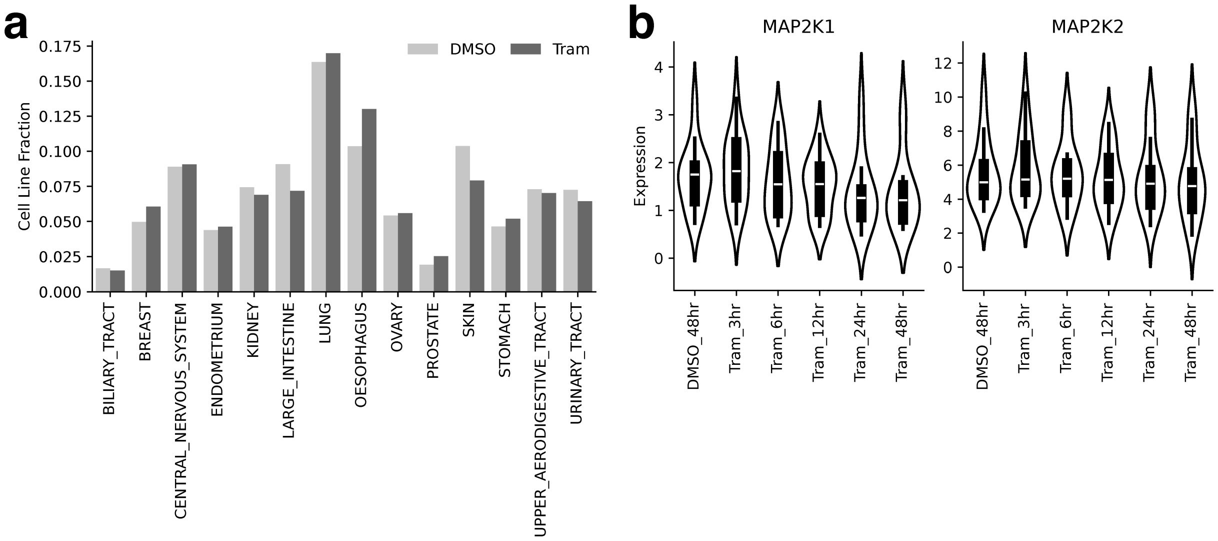


Figure S10: Cell line and expression distributions in cancer drug perturbation data[15] **a.** Cell line sample counts for dimethyl sulfoxide (DMSO), and trametinib (Tram). **b.** Distribution of expressions for primary trametinib gene targets over the course of treatment.

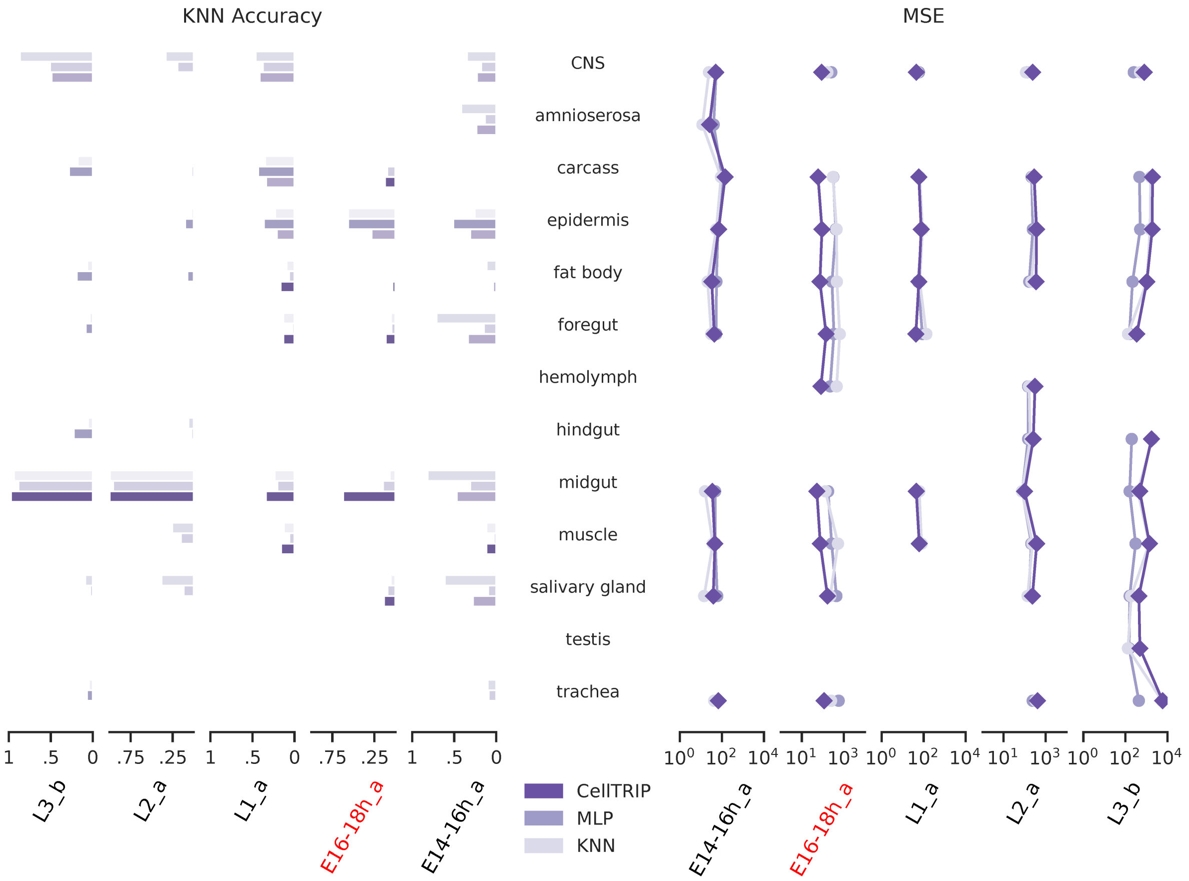


Figure S11: Performance of CellTRIP spatial imputation on *Drosophila* embryonic and larval stages Wang et al.[18]. KNN classification accuracy (Left) and MSE (Right) are shown for CellTRIP, MLP, and KNN spatial imputation methods. Results are segmented by developmental stage (x-axis) and tissue type (y-axis).
